# Remote sensing for estimating genetic parameters of biomass accumulation and modeling stability of growth curves in alfalfa

**DOI:** 10.1101/2024.04.08.588572

**Authors:** Ranjita Thapa, Karl H. Kunze, Julie Hansen, Christopher Pierce, Virginia Moore, Ian Ray, Liam Wickes-Do, Nicolas Morales, Felipe Sabadin, Nicholas Santantonio, Michael A Gore, Kelly Robbins

## Abstract

Multi-spectral imaging (MSI) collection by unoccupied aerial vehicles (UAV) is an important tool to measure growth of forage crops. Information from estimated growth curves can be used to infer harvest biomass and to gain insights in the relationship of growth dynamics and harvest biomass stability across cuttings and years. In this study, we used MSI to evaluate Alfalfa (*Medicago sativa* L. subsp. *sativa*) to understand the longitudinal relationship between vegetative indices (VIs) and forage/biomass, as well as evaluation of irrigation treatments and genotype by environment interactions (GEI) of different alfalfa cultivars. Alfalfa is a widely cultivated perennial forage crop grown for high yield, nutritious forage quality for feed rations, tolerance to abiotic stress, and nitrogen fixation properties in crop rotations. The direct relationship between biomass and VIs such as Normalized difference vegetation index (NDVI), green normalized difference vegetation index (GNDVI), red edge normalized difference vegetation index (NDRE), and Near infrared (NIR) provide a non-destructive and high throughput approach to measure biomass accumulation over subsequent alfalfa harvests. In this study, we aimed to estimate the genetic parameters of alfalfa VIs and utilize longitudinal modeling of VIs over growing seasons to identify potential relationships between stability in growth parameters and cultivar stability for alfalfa biomass yield across cuttings and years. We found VIs of GNDVI, NDRE, NDVI, NIR and simple ratios to be moderately heritable with median values for the field trial in Ithaca, NY to be 0.64, 0.56, 0.45, 0.45 and 0.40 respectively, Normal Irrigation (NI) trial in Leyendecker, NM to be 0.3967, 0.3813, 0.3751, 0.3239 and 0.3019 respectively, and Summer Irrigation Termination (SIT) trial in Leyendecker, NM to be of 0.11225, 0.1389, 0.1375, 0.2539 and 0.1343, respectively. Genetic correlations between NDVI and harvest biomass ranged from 0.52 - .99 in 2020 and 0.08 - .99 in 2021 in the NY trial. Genetic correlations for NI trial in NM for NDVI ranged from 0.72 - .98 in 2021 and SIT ranged from 0.34-1.0 in 2021. Genotype by genotype by interaction (GGE) biplots were used to differentiate between stable and unstable cultivars for locations NY and NM, and Random regression modeling approaches were used to estimate growth parameters for each cutting. Results showed high correspondence between stability in growth parameters and stability, or persistency, in harvest biomass across cuttings and years. In NM, the SIT trial showed more variation in growth curves due to stress conditions. The temporal growth curves derived from NDVI, NIR and Simple ratio were found to be the best phenotypic indices on studying the stability of growth parameters across different harvests. The strong correlation between VIs and biomass present opportunities for more efficient screening of cultivars, and the correlation between estimated growth parameters and harvest biomass suggest longitudinal modeling of VIs can provide insights into temporal factors influencing cultivar stability.

## Introduction

Alfalfa (*Medicago sativa* L. subsp. *sativa*) is one of the most widely cultivated perennial forage crops in the world with many desirable attributes such as high-yield capacity, good forage quality, tolerance to abiotic stresses, and ability to fix nitrogen and nutrient cycling (Annicchiarico et al. 2015; Hill et al. 1988). In the U.S. alfalfa is the fourth most widely grown crop with an estimated annual value of 11.7 billion dollars (USDA/ARS 2020). Alfalfa is allogamous and autotetraploid (2n = 4x = 32) and its cultivars are synthetic populations consisting of heterozygous plants (Annicchiarico and Pecetti 2021). The genetic gain in alfalfa has approached stagnation in the past few decades due to several factors including the perennial nature of the crop (long breeding cycles), multiple harvests per year, inability to make gain in harvest index due to harvesting of the entire crop, the high cost of phenotyping, tetrasomic inheritance, high genotype by environment interaction (G×E), and high levels of non-additive variance (Annicchiarico et al. 2015; Acharya et al. 2020). The narrow-sense heritability (*h^2^*) of biomass yield of alfalfa is as low as 0.20 – 0.30 (Annicchiarico 2015; Acharya et al. 2020; Riday and Brummer 2005) demanding extensive replications for phenotypic evaluation which further limits the size of breeding materials to be evaluated, ultimately leading to low selection efficiency. However, the ability to screen more materials will lead to higher effective selection intensities leading to improved response to selection.

In recent years, the advancement in high-throughput phenotyping systems, including multi-spectral imaging (MSI) platforms, have enabled the collection of high dimensional phenotypic data from large experiments and breeding trials. MSI provides an effective and non-destructive approach to evaluate the crop growth parameters throughout the crop growing season. A number of reflectance vegetation indices can be derived from spectral reflectance which have been efficiently used for large scale phenotyping and dynamic estimation of biomass greenness, nitrogen content, pigment composition, photosynthesis status and water content (Claudio et al. 2006; Mistele and Schmidhalter 2008; Schlemmer et al. 2005). MSI consists of a set of images acquired at narrow wavelength bands including both visible and near infrared (NIR) regions of the electromagnetic spectrum (Blasco et al. 2007; Chen et al. 2002). The Normalized Difference Vegetation Index (NDVI), estimated by considering the difference NIR and red wavelengths, is widely used to quantify biomass production. The green Normalized Difference Vegetation Index (GNDVI) is estimated by measuring the difference between NIR and green wavelengths and is used to measure photosynthetic activity. Other vegetation indices such as Normalized Difference Red-Edge (NDRE), Optimized Soil Adjusted Vegetation Index (OSAVI), Simplified Canopy Chlorophyll Content Index (SCCCI), and Visible Atmospherically Resistant Index (VARI_green_), have been used to predict grain yield but their use has been limited on quantification of crop biomass. Santana et al. (2021) evaluated the relationship between vegetation indices (VI)s obtained from multispectral imagery and leaf N content and yield-related traits in maize cultivars grown in different N levels, and found a positive relationship between NDVI, NDRE and grain yield under adequate N levels. Da Silva et al. (2020) evaluated the relationship between different VIs and soybean grain yield and verified a direct positive effect of NDVI and SAVI on grain yield of soybean. However, there are limited studies conducted on the relationship between different VIs and biomass yield of alfalfa crops, so further studies assessing the relationship between VIs and crop forage/biomass yield are needed. Identifying the cause-and-effect relationship between spectral and biomass yield provides an efficient phenotyping process in breeding programs. Genotypes with better spectral variables can be selected to achieve an efficient selection for biomass yield.

The use of MSI data could also be leveraged for monitoring crop growth over the growing season. Extensions of crop growth models have been proposed to incorporate functional relationships between the environmental variables and the phenotypic traits influencing yield and agronomic performance of elite breeding lines (Chapman et al. 2002; Chapman et al. 2003; Messina et al. 2015; Hammer et al. 2002; Chenu et al. 2009), and recent advancements in MSI have increased the scalability of collecting non-destructive phenotypes on a large number of experimental plots throughout the crop growth cycle. Collection of phenotypic data from multiple time points allows the monitoring of crop growth and development and hence, can increase the understanding of dynamic interactions of crop and environment.

The study of genotype by environment (G×E) interaction is one of the most important areas in plant breeding whereby breeders try to understand the stability and plasticity of the genotypes across different environments. In a perennial crop like alfalfa, the concept of persistence, or consistent performance across seasons in the same location, is a key trait for elite cultivar performance. While it may be viewed as a distinct concept from GxE, many of the same factors driving GxE are likely to play a role in persistence. For the purposes of this study we will use the terms stability, and instability, to encompass the concepts of GxE and persistence in harvest biomass yield. The traditional approach to study GxE and persistence relies on terminal traits such as harvest biomass yield, which lack the temporal resolution to study the driving factors leading to inconsistent performance across cuttings and growing seasons. In such scenarios, images taken throughout the production years of a stand can enable the longitudinal evaluation of a large number of breeding materials, providing insights into growth characteristics leading to the stability or instability of cultivar performance under differing conditions. Important growth parameters could be evaluated by studying the changes in (co)variance between adjacent time points and end-of-season traits. Quantitative genetic models can be built to accurately predict forage yields from MSI, especially given that the harvested product is imaged directly. However, the challenge lies on fitting parsimonious models that can accurately model the changes in covariance parameters across the growing season.

The phenotypic indices from high-throughput phenotyping (HTP) platforms are measured at multiple time points throughout the crop growing season and hence, are considered as longitudinal data. Repeatability models, multi-trait models, and random regression (RR) models are used to fit such longitudinal data. Repeatability models assume constant variance and correlation between measurements dates, which may not be true for longitudinal data collected at different time points throughout the crop growth cycle (Meyer and Hill 1997). In the case of multi-trait models, phenotypic traits measured at different time points are considered as distinct response variables for each cultivar. The number of parameters required to be estimated is directly related to number of time points. Hence, a strong correlation between consecutive measurements, large (co)variance matrix structure between measurements at different time points, and computational requirements restrict the application of a multi-trait model (MT) model (Speidel 2011; Anche et al. 2020). However, the RR model requires fewer parameters than MT models, can capture the change of a trait throughout the growth season, and does not require the assumption of constant variances and correlations between measurement time points (Meyer, 2020). RR models enable fitting of genetic and environmental effects over time (Schaeffer 2004), and hence results in higher accuracy of breeding values (BVs) compared to other statistical models. RR model also provide additional insights on temporal variation of biological and physiological processes underlying the trait of interest (Strucken et al. 2015) and these models have been widely used in different area of research including G×E (Calus and Veerkamp 2003; Oliveira et al. 2018). RR models commonly uses splines or Legendre polynomials to model the (co)variance of measurements at or between each time points. The objectives of this study were to (1) identify predictive image features for modeling growth and development curves for alfalfa.; (2) determine the heritability and genetic variation for image features collected throughout the growing season and (3) estimate the relationship between observed stability for development/growth parameters and stability for alfalfa biomass yield.

## Materials and Methods

### Experimental materials and biomass phenotyping

In this study, we analyzed the data from two experimental locations, (1) Cornell University Agricultural Research Experiment Station in Ithaca, NY, and (2) the Leyendecker Plant Science Research Center of New Mexico State University (NMSU) located near Las Cruces, New Mexico. A total of 36 cultivars were evaluated in the NY trial, representing both publicly released cultivars and breeding populations including ‘Guardsman II’(Viands et al., 2005), ‘Regen’ (Viands et al., 2007), ‘Algonquin’ (Baenziger, 1975),’Oneida VR’ (Viands et al., 1990), ‘Oneida Ultra’ (Viands et al., 2004), and ‘Ezra’ (Viands et al., 2012). Entries were planted on June 12, 2019, in a replicated trial with five replications in a randomized complete block design (RCBD). Plots were 6 rows of alfalfa that were 1 m by 4 m and the space between adjacent plots was 0.3 m. Forage yield was measured using a plot flail harvester, and dry matter yield for each plot was calculated from fresh forage weight and dry matter content samples. Forage yield (FY) was collected on June 5, July 9, and August 26 of 2020 and June 16, July 26, and September 13 of 2021.

A total of 24 cultivars and breeding populations with one covariate cultivar were planted in the NMSU trial on September 27, 2019. The experiment was conducted under two irrigation treatment conditions including normal irrigation (NI) and summer irrigation termination (SIT). The NI treatment received flood irrigations approximately every 14 days from March through late October. The SIT treatment only received flood irrigations from March through June and again from late September through October. Both treatment fields were planted as RCBDs with each having four replications. All experimental plots were located adjacent to a covariate plot of the cultivar, ‘NuMex Bill Melton’ (Ray et al., 2012). Each plot was comprised of three rows of alfalfa, 3.35 m in length, with 30 cm spacing between rows within a plot, and 60 cm spacings between neighboring plots and alfalfa borders. Forage biomass was harvested in 2020 with six and three harvests occurring in the NI and SIT treatments, respectively. In 2021, forage biomass was harvested seven times in the NI treatment and six times in the SIT treatment fields. All forage biomass was harvested using a Carter flail harvester to collect fresh plot weights. Subsamples of fresh chop forage were collected, weighed, and dried down to establish dry matter weights.

### Aerial phenotyping

#### NY trial

Aerial phenotyping for the NY trial commenced on April 6, 2020 in Ithaca, NY. A total of 56 flights were conducted throughout the crop growth season. A total of 7, 6, and 7 flights were flown before the first harvest (2020cut1), second harvest (2020cut2) and third harvest in 2020 (2020cut3) and a total of 22, 8, and 6 flights were flown before the first harvest (2021cut1), second harvest (2021cut1) and third harvest of 2021 (2021cut1). Four ground control points positioned at the four corners of the trial were measured with a Trimble RTK-GPS, which was used to geo-locate plots. A DJI Matrice 600 Pro unmanned aerial vehicle (UAV) equipped with a Micasense Rededge-MX multi-spectral camera was used for all flights. A flight plan was designed to obtain an 80% overlap in images collected at a flight speed of 2 m/s and an altitude of 20 m. Flights were conducted within 2 hours of solar noon on clear days when possible.

#### NMSU trial

Due to UAV equipment unavailability in 2020 and early 2021, aerial phenotyping commenced on June 3, 2021, during the third harvest cycle’s regrowth initiation for both the NI and SIT trials. A total of five harvests data from NI including NIcut3, NIcut4, NIcut5, NIcut6, NIcut7 and a total of four harvests from SIT trials including SITcut3, SITcut4, SITcut5, SITcut7 from 2021 were used for crop growth modelling and stability analysis. Ground control points were included near the four corners of each treatment field. The control points were placed on permanent stand mounts prior to each imagery flight. Upon installation, each stand was geo-located using an RTK-GPS. Multispectral imagery was captured using a DJI Matrice 600 Pro UAV and a MicaSense RedEdge-MX camera. All imagery was captured with 75% side overlap and 80% front overlap from a 20m altitude at 2.0 m/s. Imagery for both irrigation treatment fields was captured within the same flight cycle. Flights were conducted in mornings (10:00am – 12:00pm), within 3 hours of solar noon, while temperatures were cool enough to not affect UAV performance. Imagery capture occurred once per week, averaging five flights per harvest cycle, with the last flight occurring no more than two days prior to each biomass harvest. In total, 25 imaging flights were conducted over the NMSU alfalfa studies in 2021.

#### Image processing and index calculations

Orthomosacis were constructed using Pix4D mapping software (https://www.pix4d.com), and were subsequently uploaded into Imagebreed (www.imagebreed.org), a plot image database (Morales et al. 2020), for image processing and storage and calculation of vegetative indices (VI) at the plot level. Using these summary statistics, multiple VIs were calculated for each plot. Normalized difference vegetation indices (NDVI) were calculated from mean pixel values of near infrared (NIR) and Red bands of plot level images as:

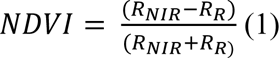

where *R_NIR_* is the near infrared reflectance and *R_R_* is the red reflectance. Green normalized difference vegetation indices (GNDVI) and Normalized difference red edge indices (NDRE) were calculated using green and red edge reflectance instead of the red reflectance in Eq. 1, respectively.

A simple Ratio was calculated as:

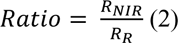

Additionally, the cumulative value of the above mentioned phenotypic indices at a specific time point, *t*, was calculated using using the rollmean function of zoo package in *R* statistical software that takes phenotypic indices values and growing degree day(s) (GDD)(s) at each time point. The equation is represented as:

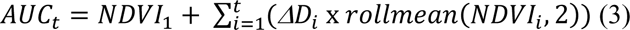

where 𝐴𝑈𝐶_𝑡_ represents the calculated AUC value at time point *t*, *Δ*𝐷_𝑖_ represents the time interval between consecutive time points (*Δ*𝐷_𝑖_ = 0 for time point 1), 𝑟𝑜𝑙𝑙𝑚𝑒𝑎𝑛(𝑁𝐷𝑉𝐼_𝑖_, 2) represents the rolling mean of NDVI values at time point *t*, and 𝑁𝐷𝑉𝐼_𝑖_ represents the NDVI value at time point *t*.

GDDs were calculated as:

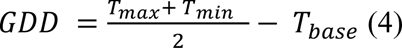

where 𝑇_𝑚𝑎𝑥_ is the maximum temperature, 𝑇_𝑚𝑖𝑛_ is the minimum temperature, and 𝑇_𝑏𝑎𝑠𝑒_ = 4 °C as the base temperature. The GDDs calculated for each time point were used as time covariates in the RR models. For the first cuttings, GDDs were calculated starting on date of planting and up to and including the date of harvest. For subsequent harvests, GDDs were calculated starting from the day after the preceding harvest. The GDDs calculated for each time point were used as time covariates in the RR models.

#### Models

A single-trait best linear unbiased prediction (ST-BLUP) model was fit to estimate the genetic and residual variances. The ST-BLUP is defined as:

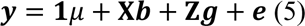

where 𝒚 is the vector of raw phenotype variables (phenotypic indices derived from MSI in this study), 𝟏 is the vector with elements of 1; 𝜇 is the overall mean; 𝒃 is the vector of fixed effect of replicate; 𝐗 is the design matrix that associates the fixed effect of replicates with response variables; 𝐙 is the design matrix with 𝒈 as a vector of random genetic effects 𝒈 ∼ 𝑁(𝟎, 𝐈𝜎 ^2^); 𝒆 is the vector of random residuals modeled as 𝒆 ∼ 𝑁(𝟎, 𝐈𝜎^2^) with an identically and independently normal distribution of residuals and 𝐈 is the identity matrix.

The ratio of estimated genetic variance to the sum of the genetic variance and residual variance was calculated to represent the broad sense heritability of biomass yield, and phenotypic indices derived from MSI.

A bi-variate multi-trait model was fit to estimate the genetic and residual correlations between biomass yield and mean values of VIs at each time point.

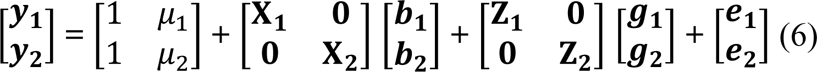

where 𝒚_𝟏_ and 𝒚_𝟐_ are the vector of response variables of traits 1 and 2; 𝜇_1_ and 𝜇_2_ are the overall means; 𝒈_𝟏_ and 𝒈_𝟐_ are the vectors of random genetic effects; 𝒃_𝟏_ and 𝒃_𝟐_ are the vectors of replication effects; 𝐗_𝟏_ and 𝐗_𝟐_ are the incidence matrices linking 𝒃_𝟏_ to 𝒚_𝟏_ and 𝒃_𝟐_ to 𝒚_𝟐_; 𝐙_𝟏_ and 𝐙_𝟐_ are the incidence matrices linking 𝒈_𝟏_ to 𝒚_𝟏_ and 𝒈_𝟐_ to 𝒚_𝟐_; 𝒆_𝟏_ and 𝒆_𝟐_ are vectors of random residual effects for trait 1 and 2, respectively. It was also assumed that [𝒈_𝟏_ 𝒈_𝟐_]∼𝑁(𝟎, 𝚺⊗ 𝐈), where 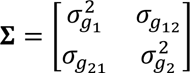 is the unstructured genetic variance and covariance matrix of the traits and 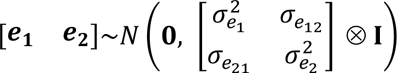.

### Random regression

Random regression models using third order of Legendre polynomials (RRLP) were used to fit a model for mean VI values and cumulative values of VI (cVI) from all time points. The biomass yield data was used as the final time point observations in the model. The variance of biomass yield was scaled to match the variance of preceding observation of VIs ensuring that yield data has similar variability pattern as VIs. The RR models were used to continuously model the (co)variance of VI and cVI measurements at different time points as a function of time.

The general random regression model for a single trait can be formulated as (Schaeffer 2004):

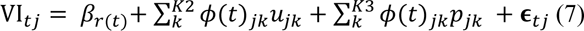

where, VI_𝑡𝑗_ is the plot level value of the *j*^th^ accession for VI at time point *t*; 𝜙(𝑡)_𝑗𝑘_ is a time covariate coefficient defined by a basis function evaluated at time point *t*; 𝛽_𝑟(𝑡)_ is the fixed effect or replicate *r* nested in time point *t*; 𝑢_𝑗𝑘_ is a *k*^th^ random regression coefficient associated with the genetic effects of the *j^th^* accession; 𝐾1 is the number of random regression parameters for fixed effect time trajectories; 𝐾2 and 𝐾3 are the number of random regression parameters for random effects; 𝑝_𝑗𝑘_ is a *k*^th^ permanent environmental random regression coefficient for the accession *j*; 𝛜_𝑡𝑗_ is the vector of residuals. The random effects at any time point were calculated as a function of the estimated RR coefficients and standardized measure of GDDs calculated from equation 3 on a per harvest basis during the growing season.

### GGE biplot analysis

The genotype main effect plus genotype by environment (GGE) biplot analysis was performed using the statistical R package called “metan” (Olivoto and Lúcio 2020). Mean biomass yield and its stability for all genotypes were visualized using GGE biplot. The GGE biplots were constructed by plotting the first principal component (PC1) against the second principal component (PC2) of the genotypes and environment calculated from a genotype-focused singular value decomposition. The following GGE biplot model was used (Yan and Kang 2002):

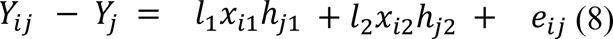

where 𝑌_𝑖𝑗_ is the mean biomass yield of genotype *i* in environment *j*; 𝑌_𝑗_ is the mean yield across all genotypes in environment *j*; 𝑙_1_ and 𝑙_2_ are the singular values for PC1 and PC2, respectively; 𝑥_𝑖1_ and 𝑥_𝑖2_ are the PC1 and PC2 scores, respectively, for genotype *i*; ℎ_𝑗1_ and ℎ_𝑗2_ are the PC1 and PC2 scores, respectively, for environment *j*; and e*_ij_* is the residual of the model associated with genotype *i* in environment *j*.

### Correlation between variance in biomass yield across environments and variance in VIs across environments

In order to have both the yield data and VIs in the same scale, z-score normalization was done by subtracting the mean (μ) from the distribution and by dividing with the standard deviation of the distribution (σ). The z-score normalization was done for each environment. Each environment was defined as a specific planting location and growth period. The correlation between the variance in yield and the variance in VIs of each genotype across locations was calculated using Pearson’s correlation method for both NY and NM trials.

## Results

### Heritability of phenotypic indices and biomass yield

For the Helfer trial, the minimum heritability of GNDVI, NDVI, NDRE, NIR and Ratio was 0, whereas the maximum heritability of GNDVI, NDVI, NDRE, NIR and Ratio was 0.92, 0.84, 0.92, 0.88 and 0.85, respectively (Fig. 1). The maximum heritability value of GNDVI and NDRE was highest among all indices followed by NIR. The median value of heritability was highest for GNDVI, followed by NDRE, NDVI, NIR and Ratio, 0.64, 0.56, 0.45, 0.45 and 0.40, respectively (Fig. 1). For 2020, the heritability of biomass yield was highest for the first harvest (0.56) followed by the third harvest (0.32) and second harvest (0.31). For 2021, the heritability was highest for the third harvest (0.62) followed by the second harvest (0.57) and the first harvest (0.31).

**Fig 1.**
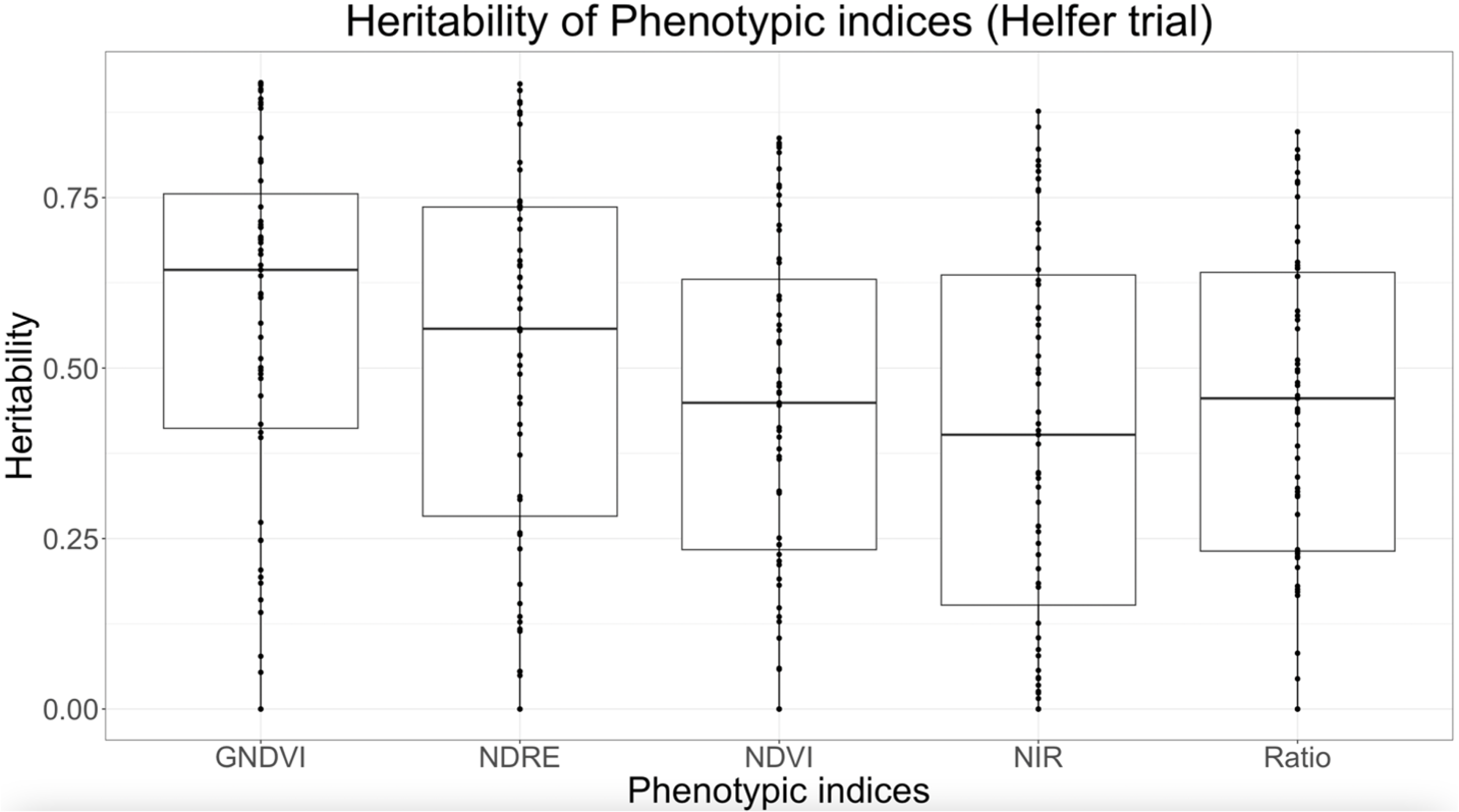
Heritability of Phenotypic indices in the Ithaca, NY trial.

For the NMSU trial in 2021, the minimum and median heritability values of the phenotypic indices under NI were higher than those under SIT whereas the maximum heritability of the phenotypic indices were higher under SIT. Under the NI, GNDVI, NDVI, NDRE, NIR and Ratio had minimum heritability values of 0.1827, 0.1076, 0.1867, 0 and 0.1112, respectively. Maximum heritability values for NMSU, GNDVI, NDVI, NDRE, NIR and Ratio were 0.7122, 0.7015, 0.6987, 0.662 and 0.6972, respectively; and median heritability values were 0.3967, 0.3813, 0.3751, 0.3239 and 0.3019, respectively (Fig. 2(a)).

**Fig 2.**
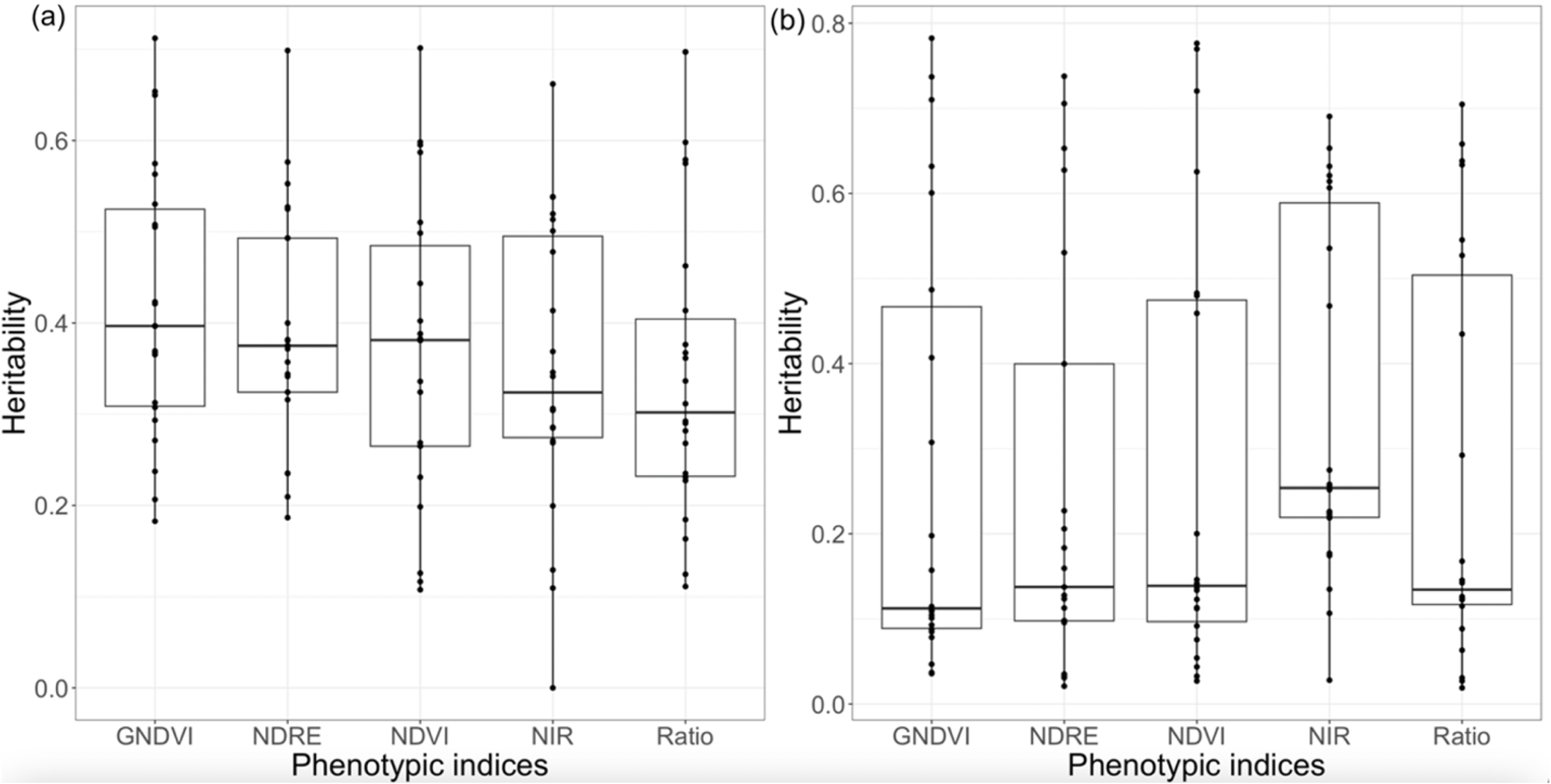
Heritability of Phenotypic indices in (a) Normal Irrigation (NMSU trial), (b) Summer termination (NMSU trial)

Under SIT, GNDVI, NDVI, NDRE, NIR and Ratio had minimum heritability values of 0.0357, 0.027, 0.0209, 0.028 and 0.0189 respectively. Maximum heritability values for GNDVI, NDVI, NDRE, NIR and Ratio were 0.7824, 0.7764, 0.7377, 0.6905 and 0.7047 respectively; and median heritability values were 0.11225, 0.1389, 0.1375, 0.2539 and 0.1343 respectively (Fig. 2(b)). Under NI, the heritability of biomass yield was highest for seventh (0.40) followed by third (0.31) and fourth (0.29). Under SIT, the heritability of biomass yield was highest for sixth (0.79) followed by the third (0.196) harvest (Fig. 2(b)).

### Phenotypic correlation of phenotypic indices and biomass yield

For the Ithaca, NY trial, the last imaging of the crop growing season was taken 9, 3 and 4 days before the first, second and third harvest of 2020, respectively, and 6, 3, and 3 days before first, second and third harvest of 2021, respectively. For both years the genetic correlation of all phenotypic indices with biomass yield was strongest for the second harvest followed by the third harvest and first harvest (Fig. 3(a), Fig. 3(b)).

**Fig 3.**
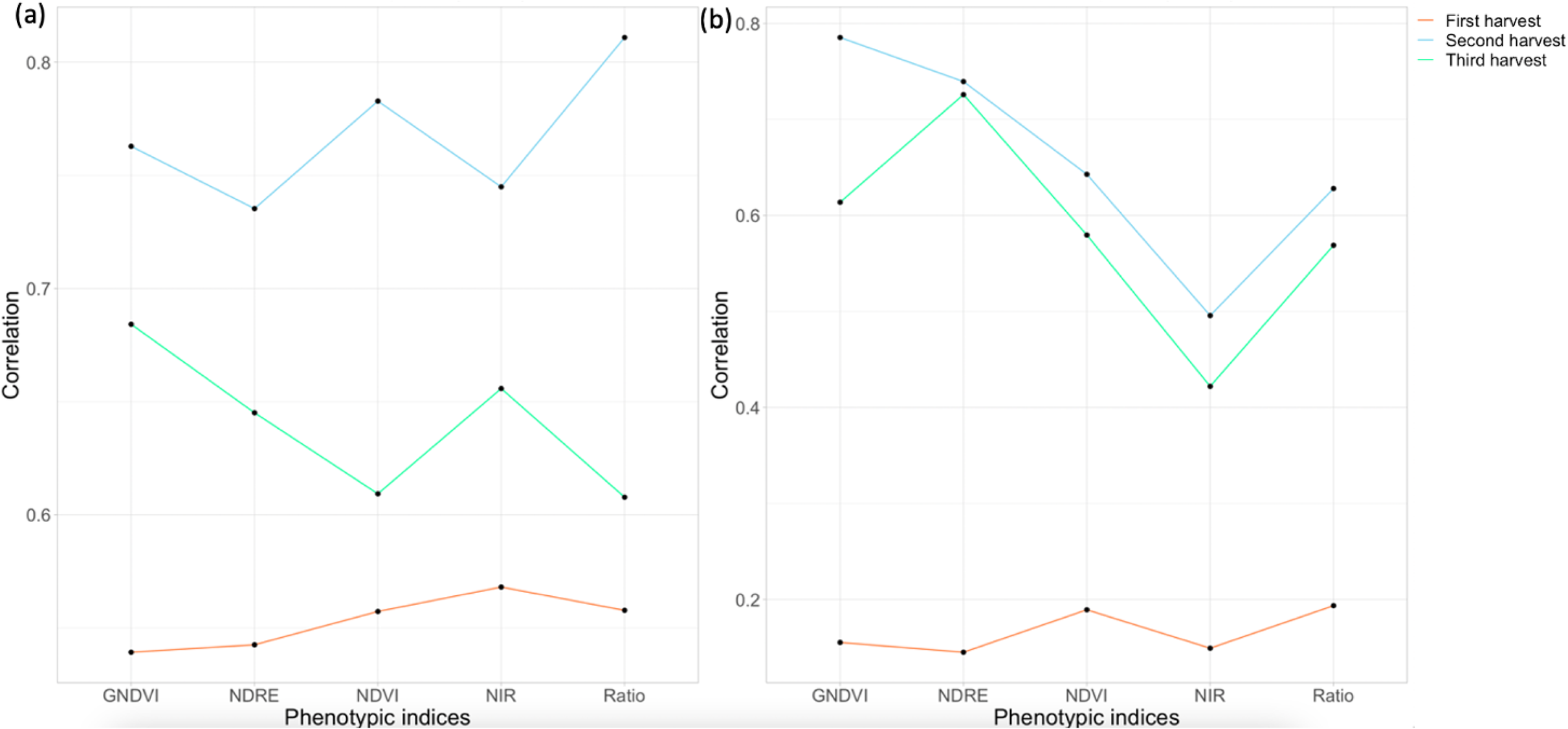
Phenotypic correlation of different phenotypic indices with biomass yield (a) Harvest year 2020 and (b) Harvest year 2021

Among all phenotypic indices in 2020, the phenotypic correlation with biomass yield was strongest for NIR (0.57) followed by Ratio (0.56) and NDVI (0.56) for the first harvest; Ratio (0.81) followed by NDVI (0.78) and GNDVI (0.76) for the second harvest; and GNDVI (0.68) followed by NIR (0.65) and NDRE (0.64) for the third harvest (Fig. 3(a)). In 2021, the phenotypic correlation with biomass yield was strongest for Ratio (0.19), followed by GNDVI (0.18) and NDVI (0.15) for the first harvest; the phenotypic correlation with biomass yield was strongest for GNDVI (0.78) followed by NDRE (0.74) and NDVI (0.64) for the second harvest; the phenotypic correlation with biomass yield was highest strongest for NDRE (0.73) followed by GNDVI (0.61) and NDVI (0.58) for the third harvest (Fig. 3(b)).

### Genetic correlation between biomass yield and phenotypic indices at different imaging time points

#### NY trial

For the first harvest of 2020, biomass yield demonstrated the highest genetic correlations with NDVI (range: 0.90 – 0.99) and NIR (range: 0.93 – 0.99) whereas biomass yield had lowest correlation with Ratio (range: 0.69 - 0.96) (Fig. 4). For the second and third harvests of 2020, Ratio showed the highest genetic correlations with ranges of 0.94 - 0.99 and 0.69 - 0.99, respectively, while NDRE had the lowest genetic correlations ranging from 0.18 to 0.94 and 0.70 to 0.98, respectively (Fig. 4).

**Fig 4.**
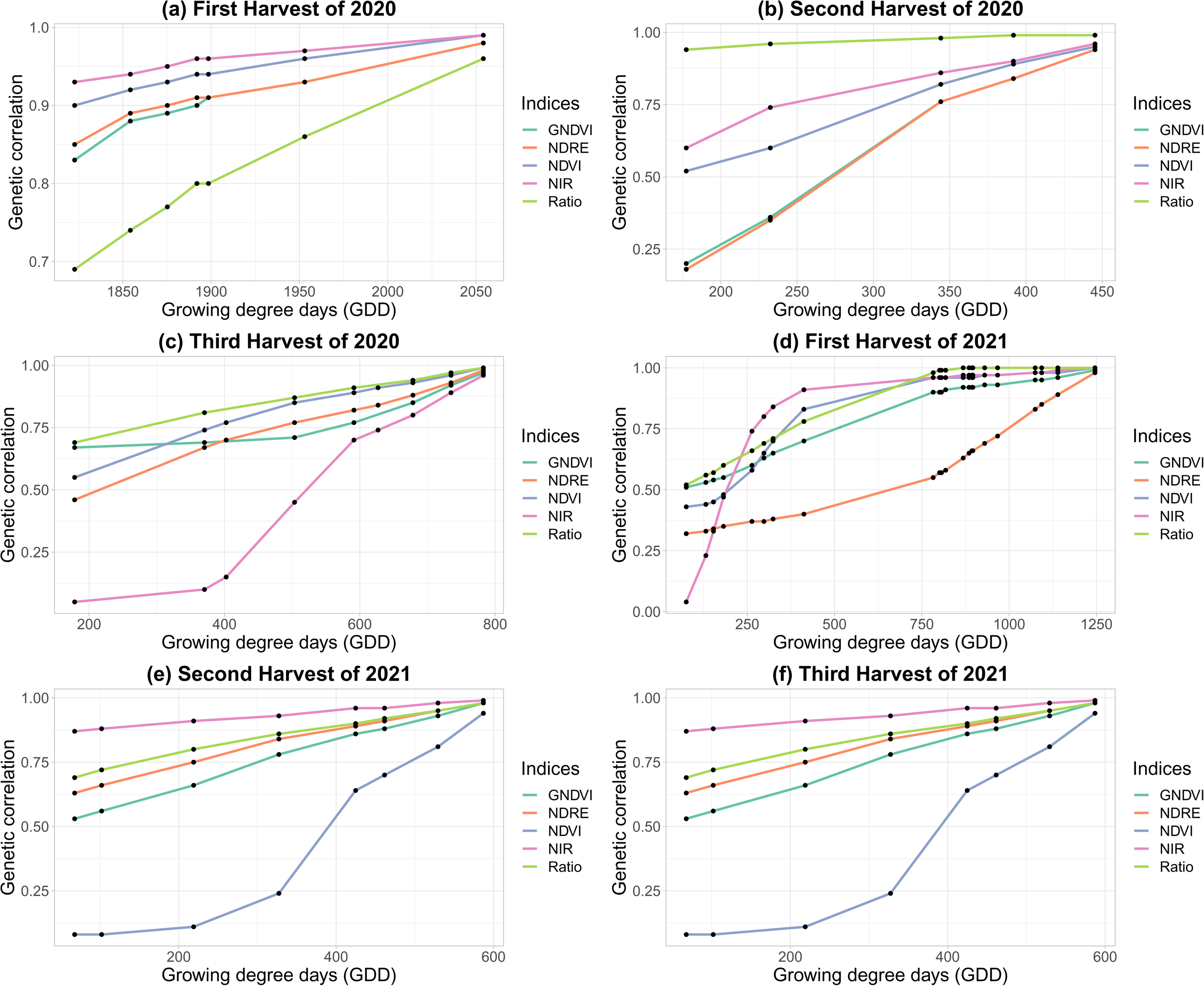
Genetic correlation of different phenotypic indices with harvest biomass yield for Ithaca, NY trial (“Helfer” field). X-axis represents Growing degree days (GDD) and Y-axis represents genetic correlation.

For the first harvest of 2021, the genetic correlation of Ratio and NIR with the biomass yield was strongest ranging from 0.68 – 0.99 and 0.1 – 0.99 respectively. The genetic correlation of NIR with biomass yield was lower than other phenotypic indices in early crop growth stage for the same harvest (Fig. 4). This pattern was only seen for one harvest out of six harvests. For the second and third harvest of 2021, the genetic correlation of NIR was strongest for second harvest and third harvest ranging from 0.84 – 0.99 and 0.91 to 1 respectively whereas genetic correlation of NDVI and GNDVI had lowest genetic correlations for second and third harvest. The genetic correlation of second and third harvest of NDVI ranged from 0.08 to 0.94 and 0.72 to 0.99 respectively for second and third harvest and the genetic correlation of second and third harvest of GNDVI ranged from 0.53 – 0.98 and 0.23 to 0.97 respectively (Fig. 4).

#### NMSU trial

Under NI, the genetic correlation of NDVI and Ratio at all imaging time points were highest for all harvests except for June 25 to Jul 22 regrowth cycle of 2021 (Fig. 5). The genetic correlation of NDVI ranged from 0.8 to 0.97 for May 28 to June 24 regrowth cycle, 0.72 to 0.97 for June 25 to Jul 22 regrowth cycle, 0.78 to 0.96 for July 23 to August 27 regrowth cycle, 0.77 to 0.97 for August 28 to September 29 regrowth cycle, 0.88 to 0.98 for September 30 to November 12 regrowth cycle and the genetic correlation of Ratio ranged from 0.69 to 0.95 for May 28 to June 24 regrowth cycle, 0.69 to 0.97 for June 25 to Jul 22 regrowth cycle, 0.69 to 0.94 for July 23 to August 27 regrowth cycle, 0.69 to 0.96 for August 28 to September 29 regrowth cycle, and 0.69 to 0.97 for September 30 to November 12 regrowth cycle (Fig. 5).

**Fig 5.**
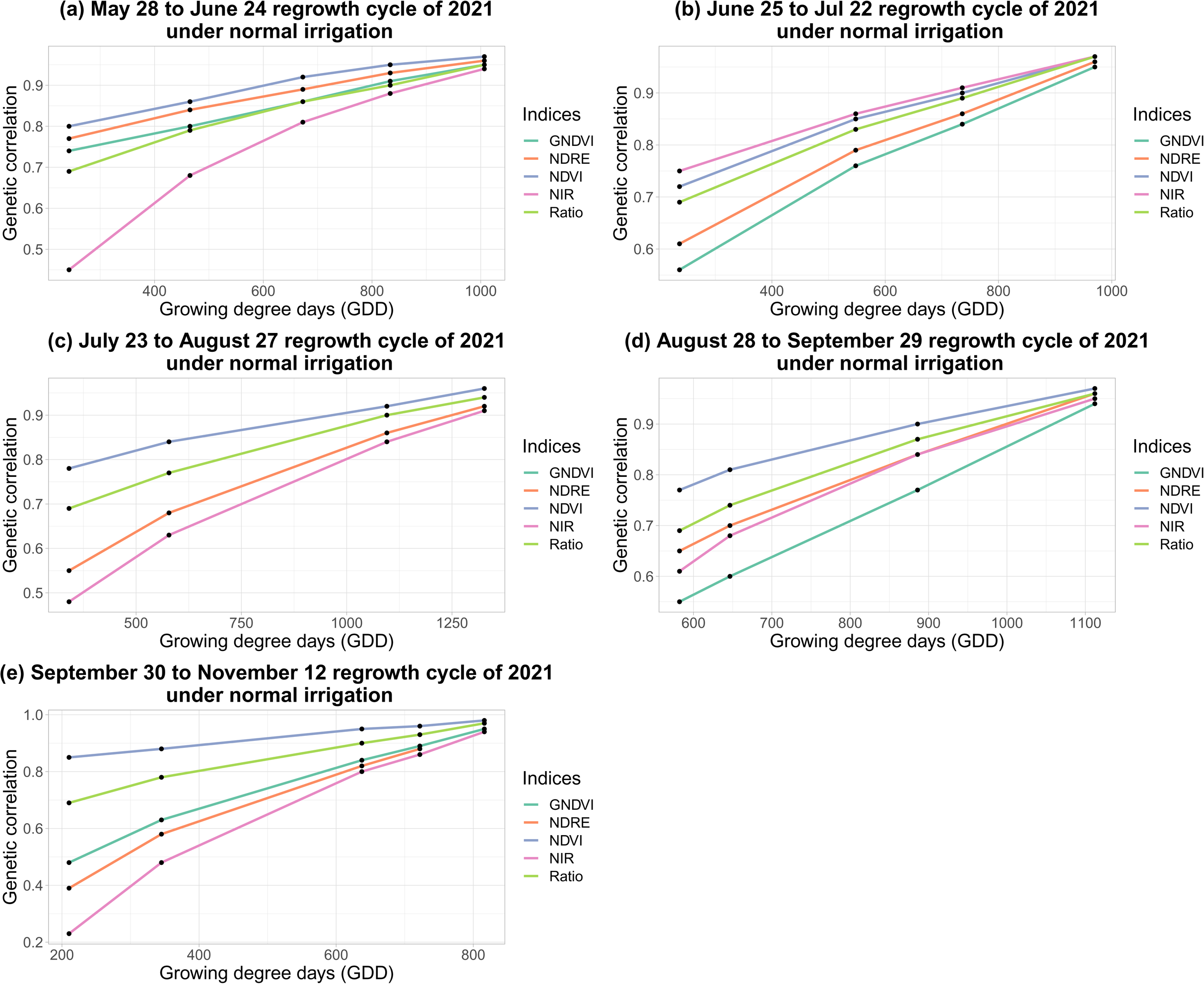
Genetic correlation of different phenotypic indices with final harvest biomass yield under normal irrigation condition of NMSU trial. X-axis represents Growing degree days (GDD) and Y-axis represents genetic correlation.

Under SIT, NDVI and Ratio had highest genetic correlation with biomass yield compared to other phenotypic indices. Genetic correlations ranged from 0.84 to 0.97 for May 28 to June 25 regrowth cycle, 0.91 to 0.97 for June 26 to Jul 22 regrowth cycle, 0.99 to 1 for July 23 to August 26 regrowth cycle and 0.69 to 0.99 for August 27 to November 11 regrowth cycle for NDVI and genetic correlation ranged from 0.69 to 0.95 for May 28 to June 25 regrowth cycle, 0.69 to 0.91 for June 26 to Jul 22 regrowth cycle, 0.69 to 1 for July 23 to August 26 regrowth cycle and 0.69 to 0.99 for August 27 to November 11 regrowth cycle for Ratio (Fig. 6).

**Fig 6.**
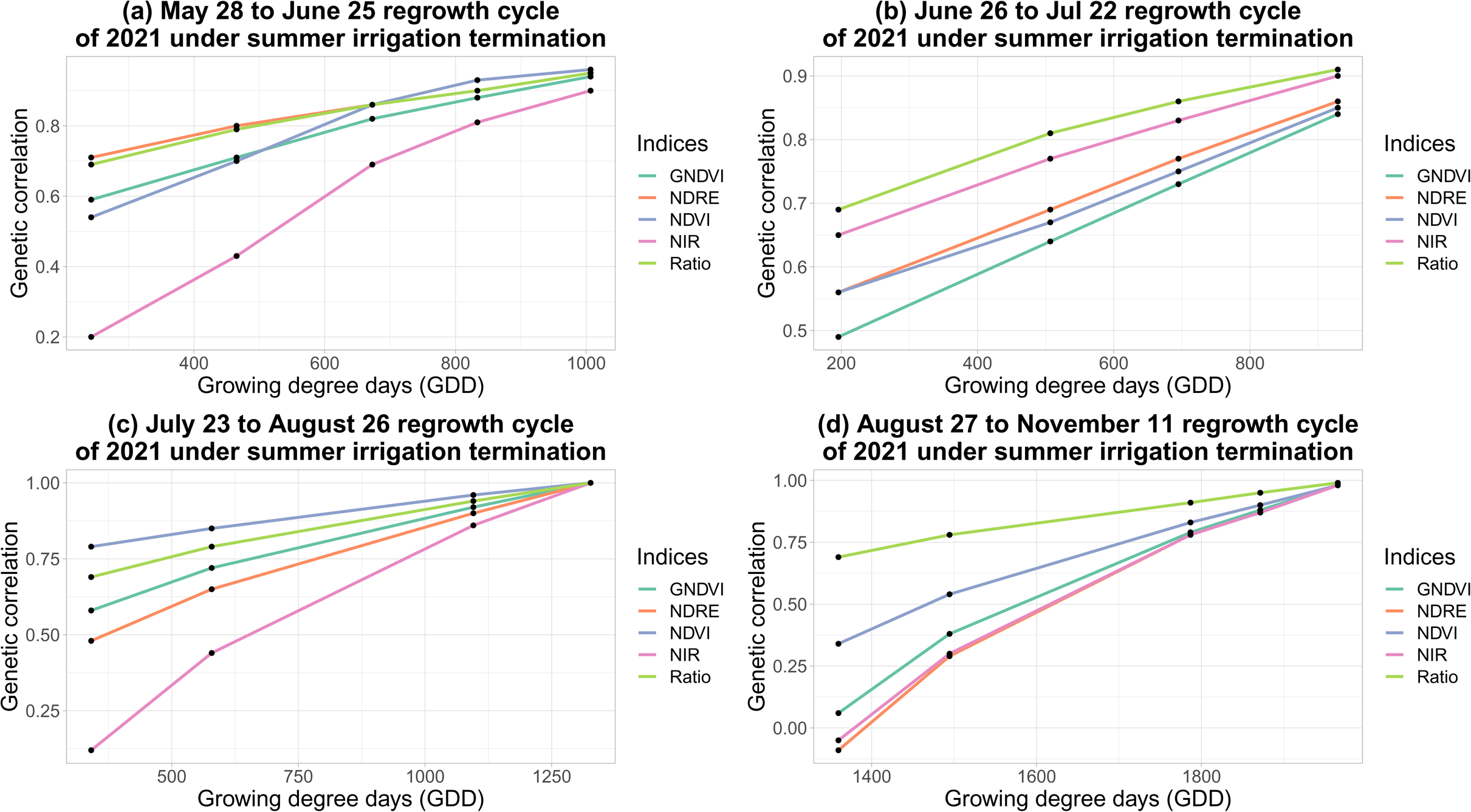
Genetic correlation of different phenotypic indices with final harvest biomass yield under summer irrigation termination condition of NMSU trial. X-axis represents Growing degree days (GDD) and Y-axis represents genetic correlation.

### Genetic correlation among phenotypic indices at different imaging time points

#### Ithaca, NY trial

The genetic correlation of phenotypic indices at different time points were evaluated running multi-trait models Supplemental Figure 1 (a) to (e)). The genetic correlation among NIR and Ratio at different time points were strongest compared to other indices (Supplemental Figure 1 (d), Supplemental Figure 1 (e)). The genetic correlation of Ratio at different time points were greater than 0.71 for all harvests of 2020 and 2021 except for the first harvest of 2021, where genetic correlations between the first time point and the last 14 time points ranged from 0.49 to 0.67. The genetic correlations among NIR at different time points were greater than 0.65 for all harvests of 2020 and 2021 except for third harvest of 2020 and first harvest of 2021, where the genetic correlations ranged from −0.17 to 0.04 between first and last five imaging time points and 0.04 to 0.05 between first and last 14 imaging time points. The genetic correlations of NDVI, GNDVI, and NDRE at different time points were in the range of 0.52 – 1, 0.2 −1, and 0.18 – 0.1, respectively, for all three harvests of 2020 and the third harvest of 2021. Genetic correlations were lower for first and second harvest of 2021 across all harvests (Supplemental Figure 1 (a) to (e)). The genetic correlation of cumulative value of all the indices from second time point to other time points were 1 whereas the genetic correlation of cumulative value of all the indices of first time point with other time points were in the range of 0.9 – 0.99 (Supplemental Figure 2 (a) to (e)).

#### NMSU trial

Under NI, among all phenotypic indices, the genetic correlation of NDVI and Ratio at different time points were strongest (Supplemental Figure 3 (c), Supplemental Figure 3 (e)). The genetic correlation of Ratio at different time points ranged from 0.69-0.98, 0.69-0.97, 0.69-0.98, 0.69-0.99, 0.69-0.98 for the May 28 to June 24 regrowth cycle, June 25 to Jul 22 regrowth cycle, July 23 to August 27 regrowth cycle, August 28 to September 29 regrowth cycle and September 30 to November 12 regrowth cycle, respectively (Supplemental Figure 3 (e)). Similarly, the genetic correlation of NDVI at different time points ranged from 0.72 - 0.99, 0.74 – 0.97, 0.76 – 0.98, 0.76 – 0.99 and 0.81 – 0.99 for the May 28 to June 24 regrowth cycle, June 25 to Jul 22 regrowth cycle, July 23 to August 27 regrowth cycle, August 28 to September 29 regrowth cycle and September 30 to November 12 regrowth cycle, respectively (Supplemental Figure 3 (c)). The genetic correlations of NIR, GNDVI, and NDRE at different time points were lowest compared to other indices (Supplemental Figure 3 (d), Supplemental Figure 3 (a), Supplemental Figure 3 (b)). The genetic correlation of the cumulative value of all the phenotypic indices of first time points with other time points were 0.99 and 1.0 for all other time points (Supplemental Figure 4 (a) to (e)).

Similarly, under SIT, the genetic correlation of Ratio and NDVI at different time points were strongest (Supplemental Figure 5 (a) to (e)). The genetic correlation of Ratio at different time points ranged from 0.69 – 0.98, 0.69 – 0.98, 0.69 – 1 and 0.69 – 0.99 for the May 28 to June 25 regrowth cycle, June 26 to Jul 22 regrowth cycle, July 23 to August 26 regrowth cycle and August 27 to November 11 regrowth cycle, respectively (Supplemental Figure 5 (e)). Similarly, the genetic correlation of NDVI ranged from 0.54 – 0.97, 0.56 – 0.97, 0.79 – 1, 0.34 – 0.98 for the May 28 to June 25 regrowth cycle, June 26 to Jul 22 regrowth cycle, July 23 to August 26 regrowth cycle and August 27 to November 11 regrowth cycle, respectively (Supplemental Figure 5 (a)). Among the other indices at different time points, the genetic correlation among GNDVI ranged from 0.59 – 0.97, 0.49 – 0.96, 0.58 – 1, 0.06 – 0.98 for the third, fourth, fifth and seventh harvest, respectively (Supplemental Figure 5 (a)), NDRE ranged from 0.71 – 0.98, 0.56 – 0.97, 0.48 – 1, −0.09 – 0.97 for the May 28 to June 25 regrowth cycle, June 26 to Jul 22 regrowth cycle, July 23 to August 26 regrowth cycle and August 27 to November 11 regrowth cycle harvest, respectively (Supplemental Figure 5 (b)), and NIR ranged from 0.2 – 0.95, 0.65 – 0.98, 0.12 – 1, and −0.05 – 0.98 respectively for May 28 to June 25 regrowth cycle, June 26 to Jul 22 regrowth cycle, July 23 to August 26 regrowth cycle and August 27 to November 11 regrowth cycle, respectively (Supplemental Figure 5 (d)). The genetic correlation of the cumulative value of all the phenotypic indices of first time points and other time points were 0.99 whereas for other time points were 1 (Supplemental Figure 6 (a) to (e)).

### Growth curve analysis using genetic merit calculated from Random Regression Model

The temporal growth curves of all alfalfa genotypes were constructed using breeding values calculated using RRLP and different phenotypic indices as longitudinal phenotypic traits (Supplemental Figures 7, 8, 9). The high-resolution temporal growth curves of different alfalfa genotypes showed clear differences between high yielding and low yielding genotypes. Both raw values of phenotypic indices and their respective cumulative values were used to run RR model. Compared to the raw value, cumulative value of phenotypic indices gave better model fit and higher resolution of temporal growth curves (Supplemental Figure 7(f) to 7(j), Supplemental Figure 9(f) to 9(j)). Using the raw value of phenotypic indices as the phenotypic trait, a larger spread in breeding values of the genotypes were observed in the early stages of growth, indicating greater genetic variance captured by the proximal sensing phenotypes in early growth stages.

**Fig. 7.**
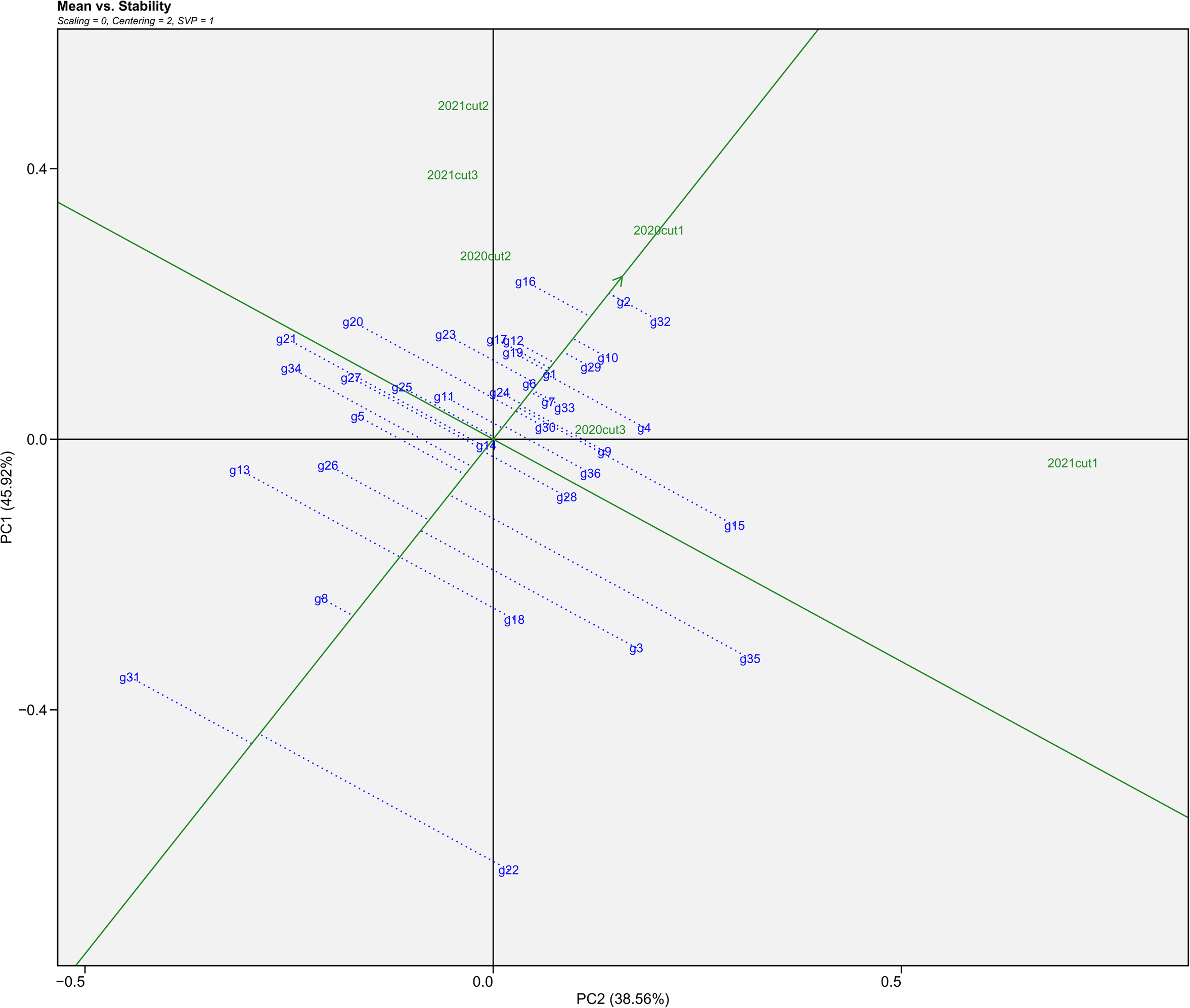
The “mean vs. stability” view of the genotype main effects plus genotype environment interaction (GGE) biplot based on genotype environment yield data of 36 alfalfa genotypes evaluated in six environments (First, Second and Third Harvest of 2020, and First, Second and Third Harvest of 2021) of Helfer field in Ithaca, NY.

### GGE biplot analysis

The GGE Biplots in Figs. 7 and 8, provide a “mean versus stability” graph of cultivar performance in NY and NMSU trials, respectively (Yan et al. 2007). The green single arrowed line, referred to as the “average environment axis”, provides an indication of the mean performance of cultivars, with the arrow pointing to a greater value according to their mean performance across all environments. The green line that is perpendicular to the average environment axis, provides an indication of stability in cultivar performance across environments. As such, cultivars with projections closer to the average environment axis exhibited more stable performance for harvested biomass across cuttings and years. An ideal cultivar would have a high mean performance, further along in the average environment axis in the direction indicated by the arrow, and would show stable performance with a projection near the average environment axis. For the cultivars tested in NY (Fig. 7), the cultivars g1, g2, g10, g29, and g32 were relatively stable and high yielding, and g8 was relatively stable and low yielding. Cultivars g3, g13, g18, g22, g31 were relatively unstable and low yielding, and g4, g15, g20, g23 were relatively unstable and high yielding. A similar analysis was applied to the NMSU trial data (Fig. 8), identifying G13, G14 and G15 as stable and low yielding cultivars, and G25, G7 and G9 as relatively stable and high yielding cultivars. Results indicate that G24, G23 and G21 were relatively unstable and high yielding, and G2, G8, G15, G17 were relatively unstable and low yielding.

**Fig. 8.**
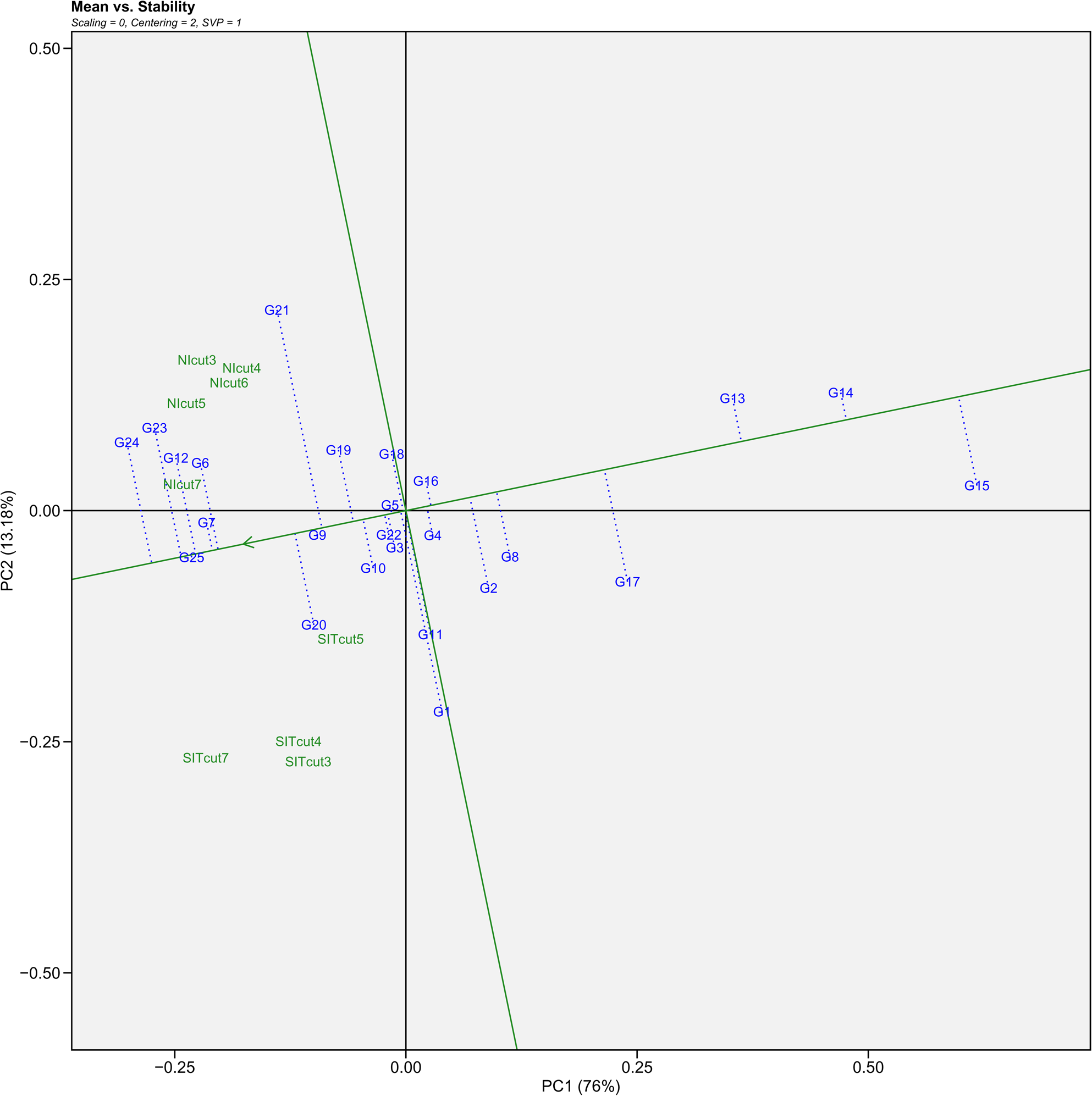
The “mean vs. stability” view of the genotype main effects plus genotype environment interaction (GGE) biplot based on genotype environment yield data of 24 alfalfa genotypes and one covariate (G4) evaluated in nine environments - NIcut3, NIcut4, NIcut5, NIcut6, and NIcut7 of normal irrigation and SITcut3, SITcut4, SITcut5 and SITcut7 of summer irrigation termination of NMSU.

**Fig. 9.**
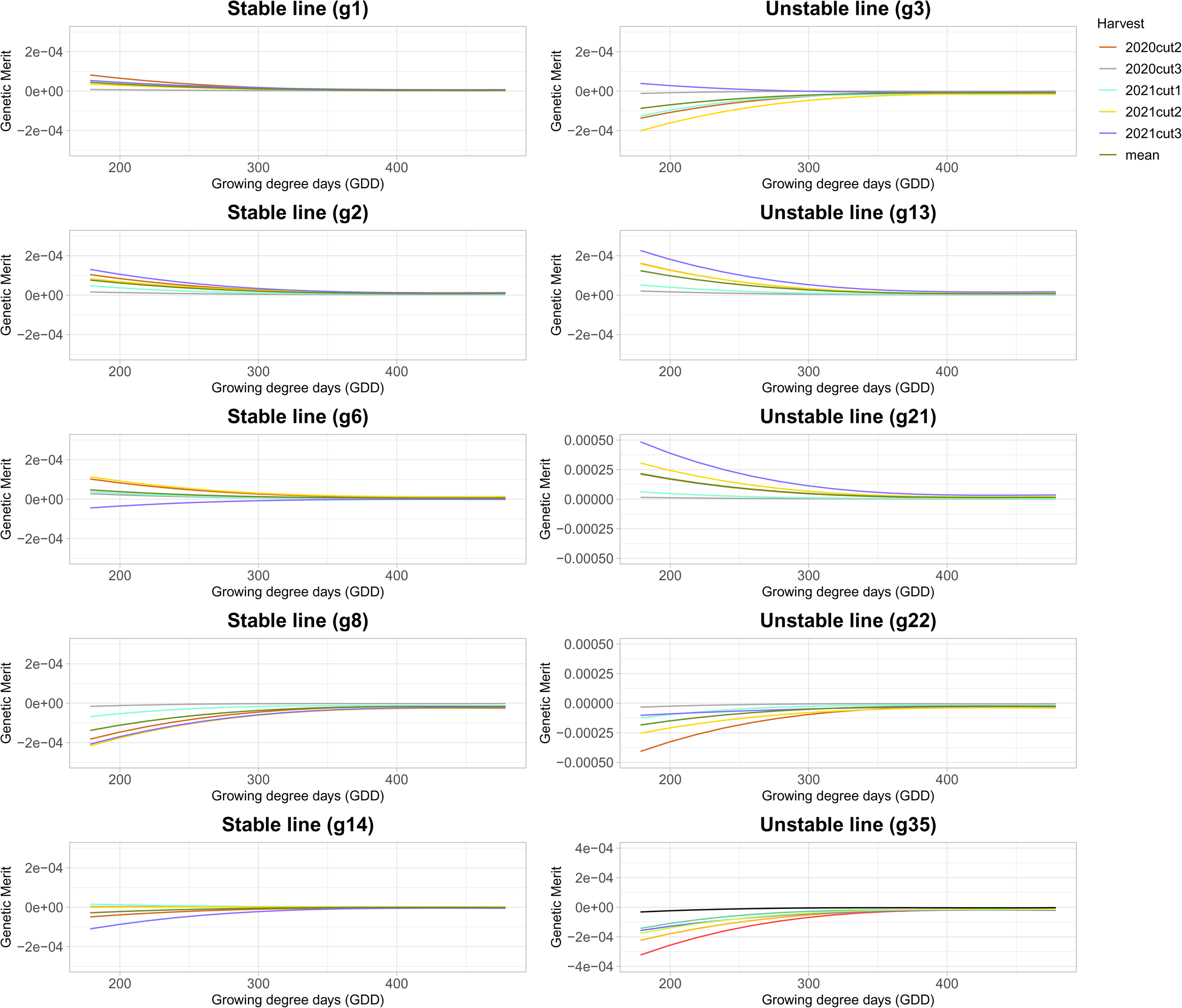
Growth curves derived from GNDVI of five stable and five unstable alfalfa cultivars across five different harvest seasons (excluding first harvest) of the Ithaca, NY trial. X-axis indicates Growing degree days (GDD) and Y-axis indicates breeding values estimated using Random Regression model with third order of Legendre polynomials.

### Stability and plasticity analysis using a growth curve modeling approach

Among the most unstable and stable genotypes identified from GGE biplot analysis, five stable and five unstable cultivars were selected (Fig.7, Fig.8). To determine whether stability in biomass yield across cuttings was reflected in the plasticity of the plant growth, the growth curves of these most stable and unstable cultivars across different environments were plotted (Fig.9 - Fig.25). Results showed high variance in the growth curves of unstable cultivars across all cuttings when compared to the stable cultivars in the Ithaca, NY trial (Fig.9 - Fig.13). Similar, although less pronounced, results were observed for NMSU trial (Fig.14 - Fig.17). Among all phenotypic indices, the growth curves estimated using NDVI and Ratio at Ithaca, NY were found to give clear separation in the stability and plasticity (Fig.10, Fig.13). Both the stable and unstable cultivars were found to be more stable in NI than SIT of NMSU (Fig.18 – Fig. 25), with differing growth patterns between the two irrigation treatments.

**Fig. 10.**
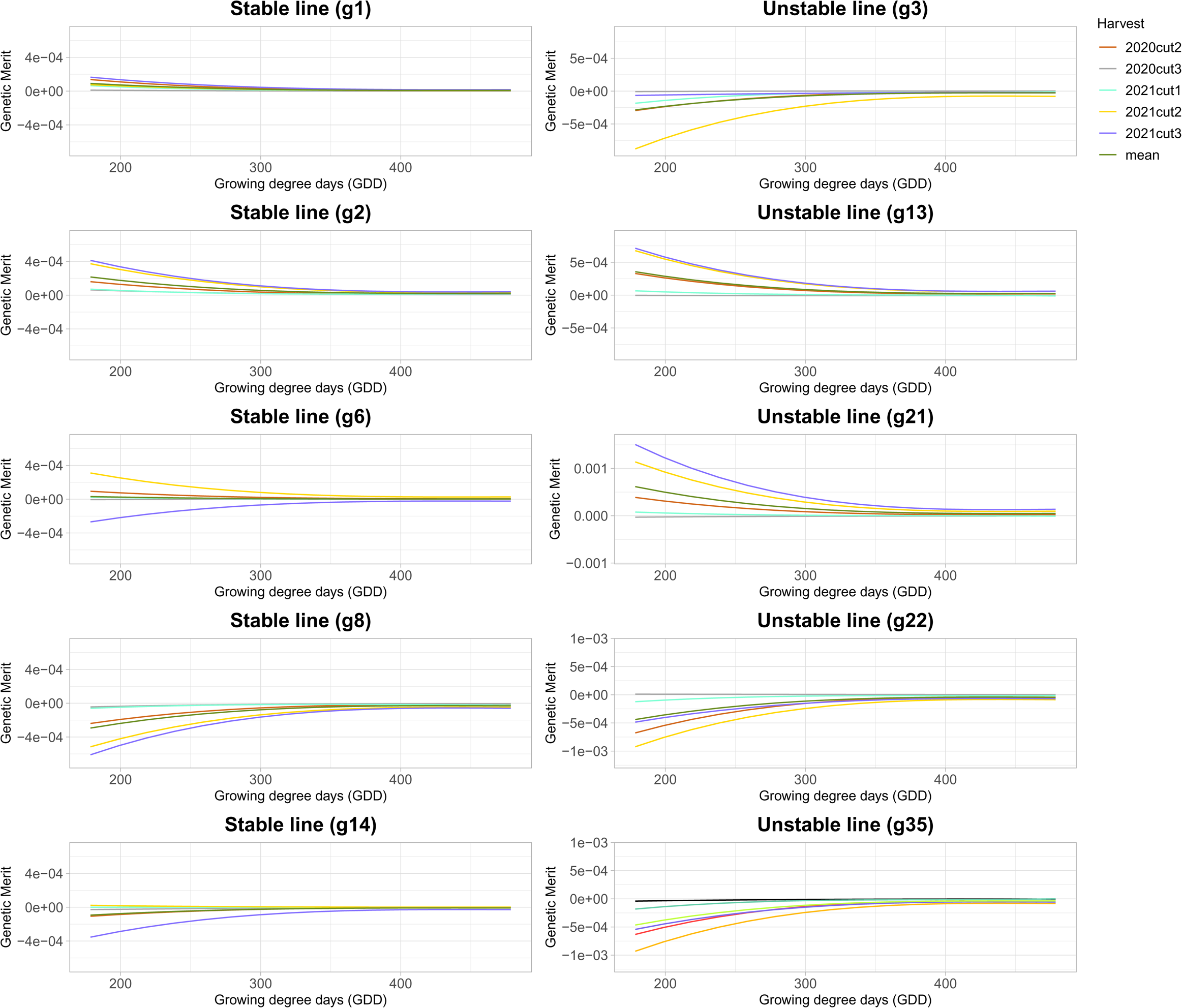
Growth curves derived from NDVI of five stable and five unstable alfalfa cultivars across five different harvest seasons (excluding first harvest) of the Ithaca, NY trial. X-axis indicates Growing degree days (GDD) and Y-axis indicates breeding values estimated using Random Regression model with third order of Legendre polynomials.

**Fig. 11.**
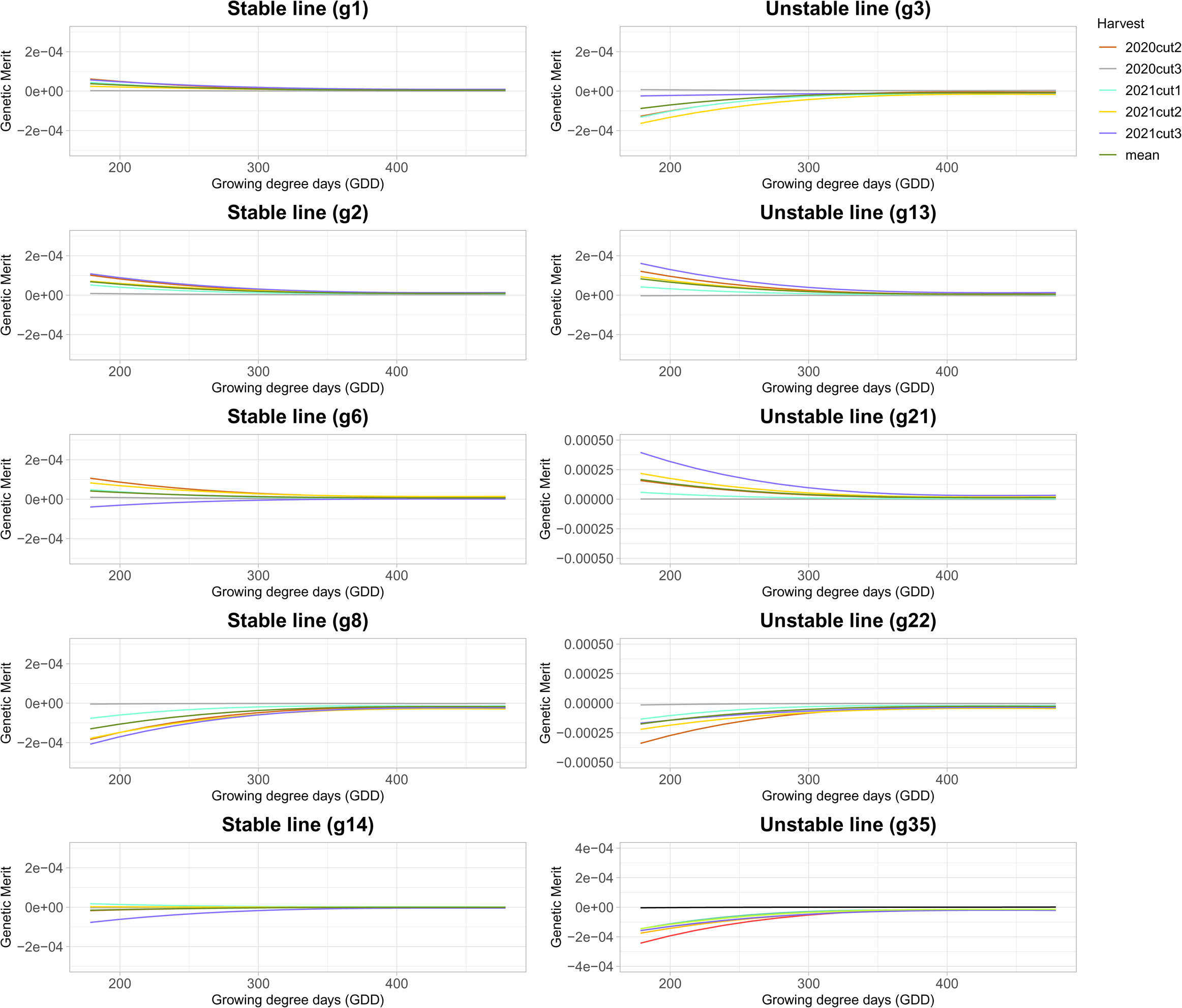
Growth curves derived from NDRE of five stable and five unstable alfalfa cultivars across five different harvest seasons (excluding first harvest) of the Ithaca, NY trial. X-axis indicates Growing degree days (GDD) and Y-axis indicates breeding values estimated using Random Regression model with third order of Legendre polynomials.

**Fig. 12.**
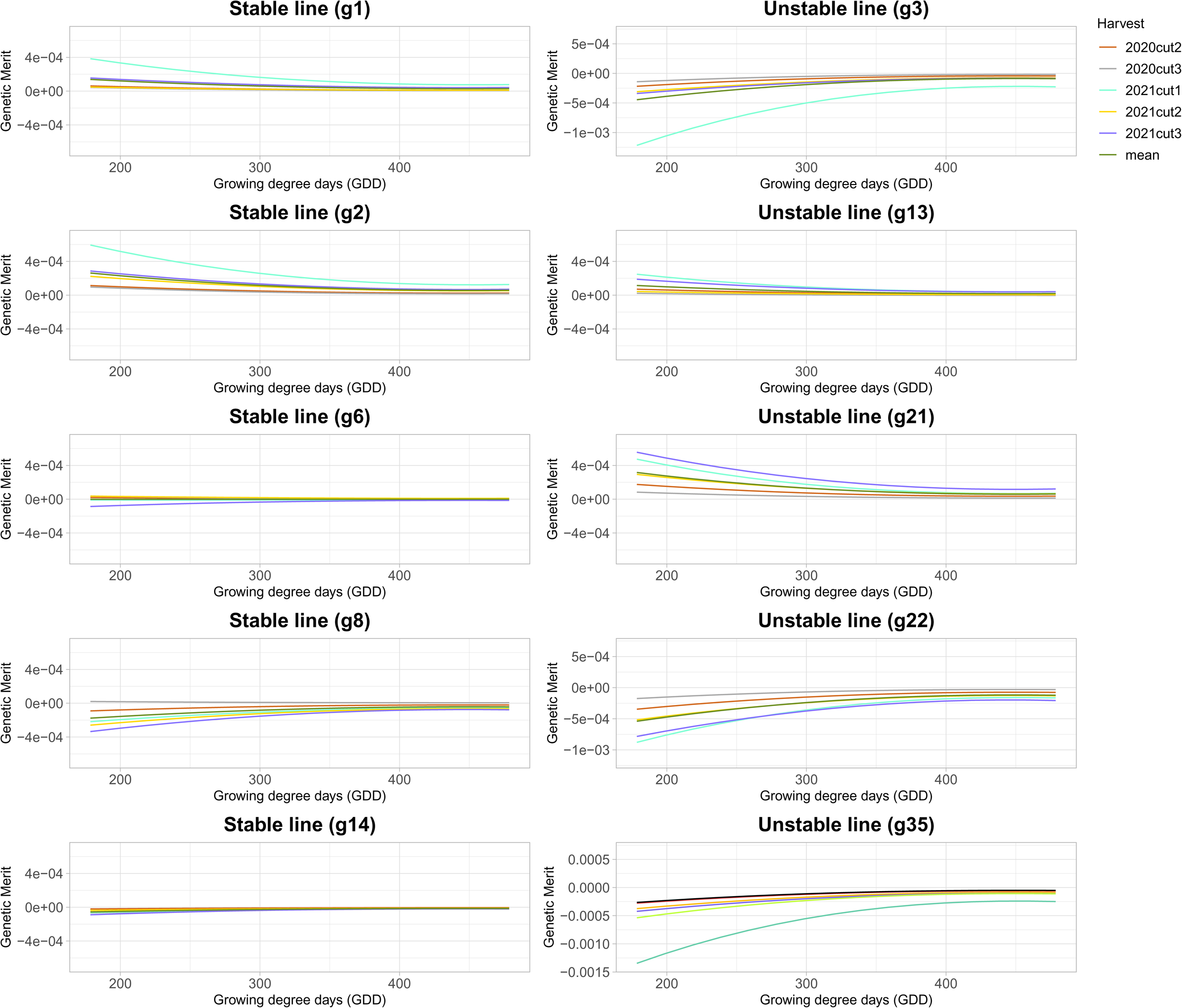
Growth curves derived from NIR of five stable and five unstable alfalfa cultivars across five different harvest seasons (excluding first harvest) of the Ithaca, NY trial. X-axis indicates Growing degree days (GDD) and Y-axis indicates breeding values estimated using Random Regression model with third order of Legendre polynomials.

**Fig. 13.**
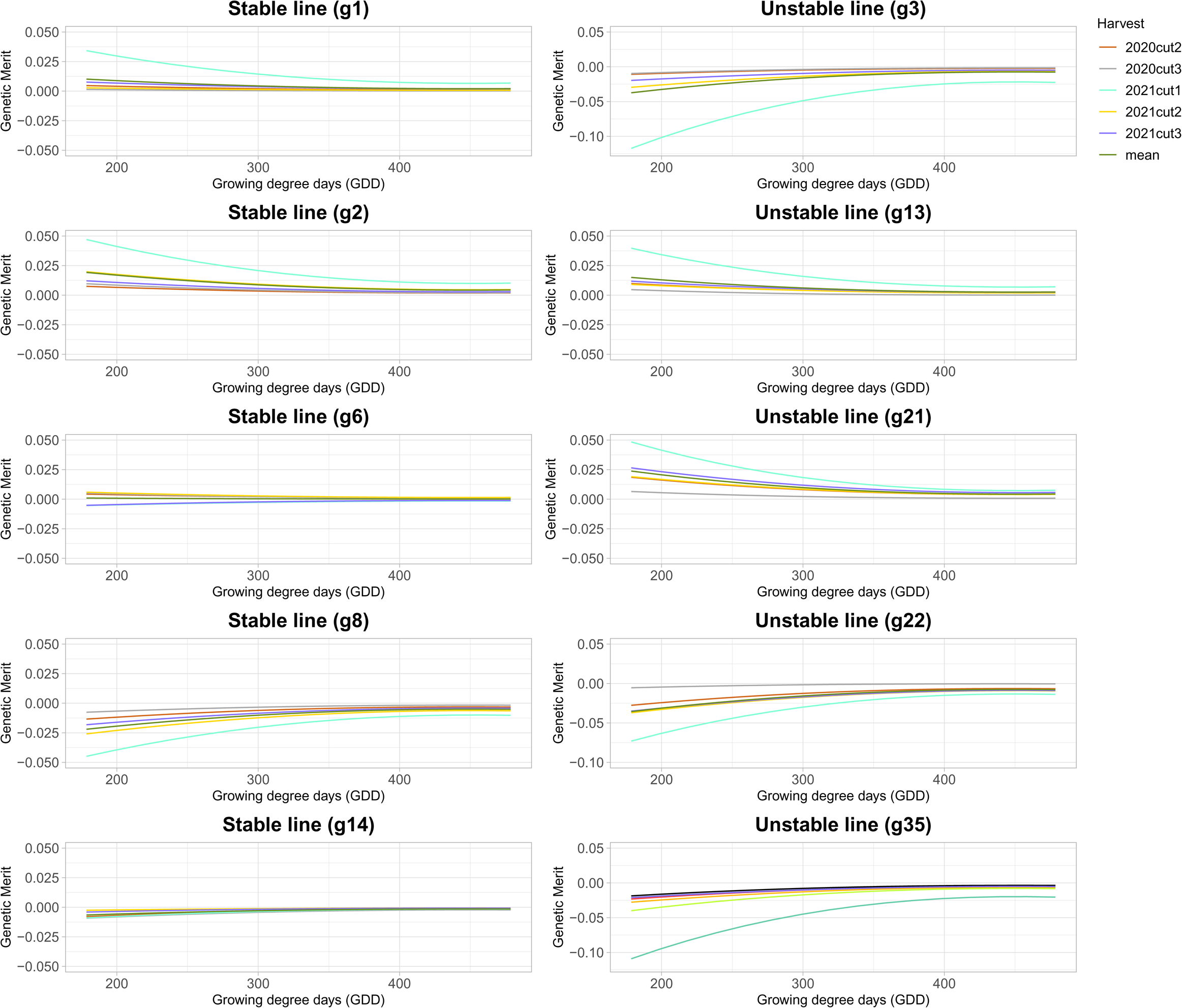
Growth curves derived from Ratio of five stable and five unstable alfalfa cultivars across five different harvest seasons (excluding first harvest) of the Ithaca, NY trial. X-axis indicates Growing degree days (GDD) and Y-axis indicates breeding values estimated using Random Regression model with third order of Legendre polynomials.

**Fig. 14.**
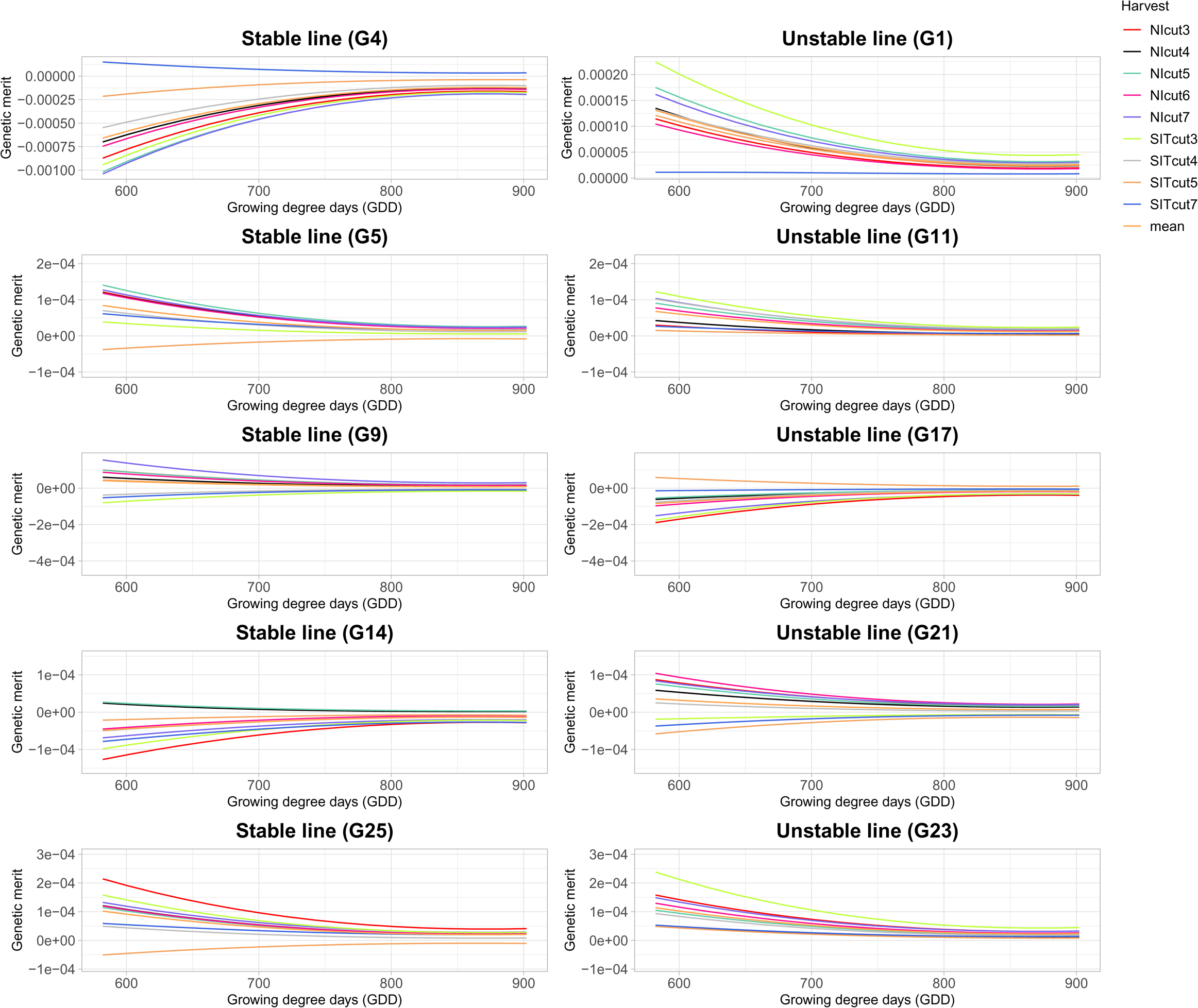
Growth curves derived from GNDVI of five stable and five unstable alfalfa cultivars across nine different harvest seasons of the NMSU trial. X-axis indicates Growing degree days (GDD) and Y-axis indicates breeding values estimated using Random Regression model with third order of Legendre polynomials.

Using GNDVI, NDVI and Ratio as the phenotypic trait, the variance of stable cultivars g1, g2, g6, g8, and g14 across all environments were less than the variance of unstable cultivars g3, g13, g21, g22, g35 (Fig. 9, 10, 13) in Ithaca, NY trial. There was more separation during early growth among the cultivars than at later timepoints. Similar results were observed for NMSU trial. The temporal growth curves derived from NDVI, NIR and Ratio were found to be the best discriminating the variance of genetic merit for stable and unstable cultivars across different harvests (Fig. 15, Fig. 16, Fig. 17).

**Fig. 15.**
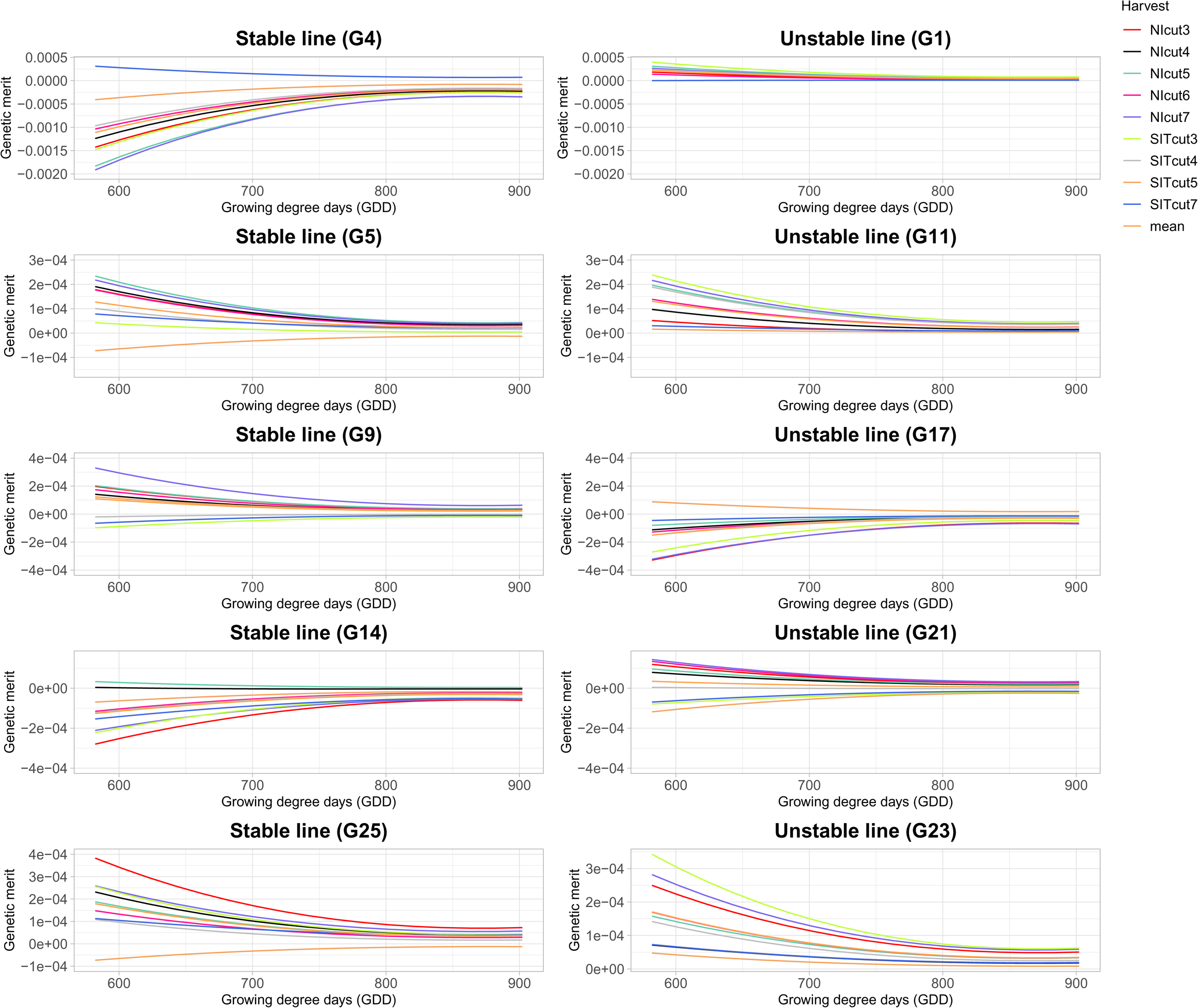
Growth curves derived from NDVI of five stable and five unstable alfalfa cultivars across nine different harvest seasons of the NMSU trial. X-axis indicates Growing degree days (GDD) and Y-axis indicates breeding values estimated using Random Regression model with third order of Legendre polynomials.

**Fig. 16.**
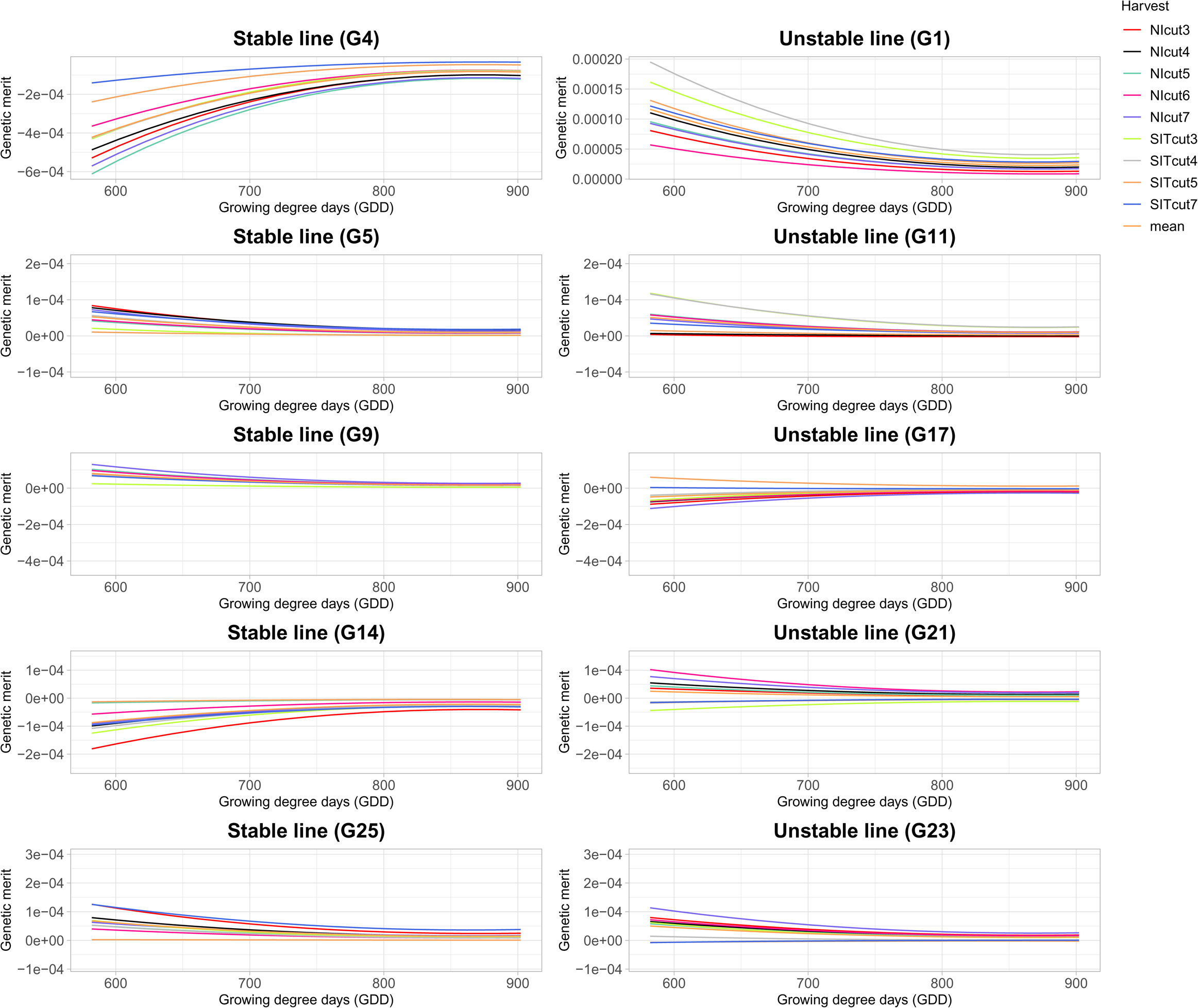
Growth curves derived from NIR of five stable and five unstable alfalfa cultivars across nine different harvest seasons of the NMSU trial. X-axis indicates Growing degree days (GDD) and Y-axis indicates breeding values estimated using Random Regression model with third order of Legendre polynomials.

**Fig. 17.**
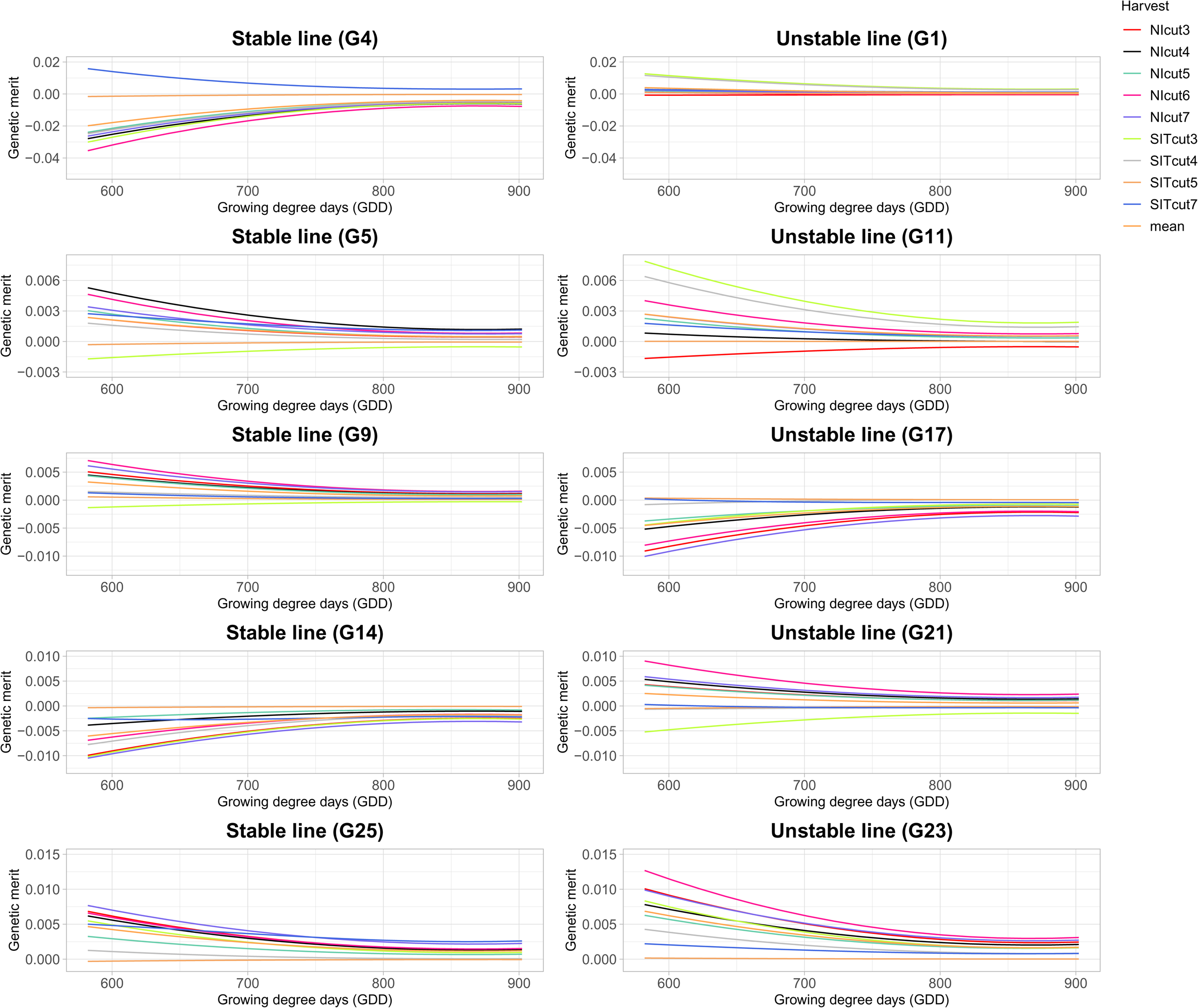
Growth curves derived from Ratio during the growing season of five stable and five unstable alfalfa cultivars across nine different harvest seasons of the NMSU trial. X-axis indicates Growing degree days (GDD) and Y-axis indicates breeding values estimated using Random Regression model with third order of Legendre polynomials.

### Stability and plasticity analysis across different irrigation conditions

The growth trajectories of stable and unstable cultivars were compared separately across all cuttings of NI and SIT conditions (Fig. 18 to Fig. 25). The variance in growth curves of both stable cultivars derived from GNDVI, NDVI and Ratio were found to be higher in summer irrigation termination condition than in normal irrigation condition (Fig. 18, Fig. 20, Fig. 24). Similar results were observed for unstable cultivars (Fig. 19, Fig. 21, Fig. 25).

**Fig. 18.**
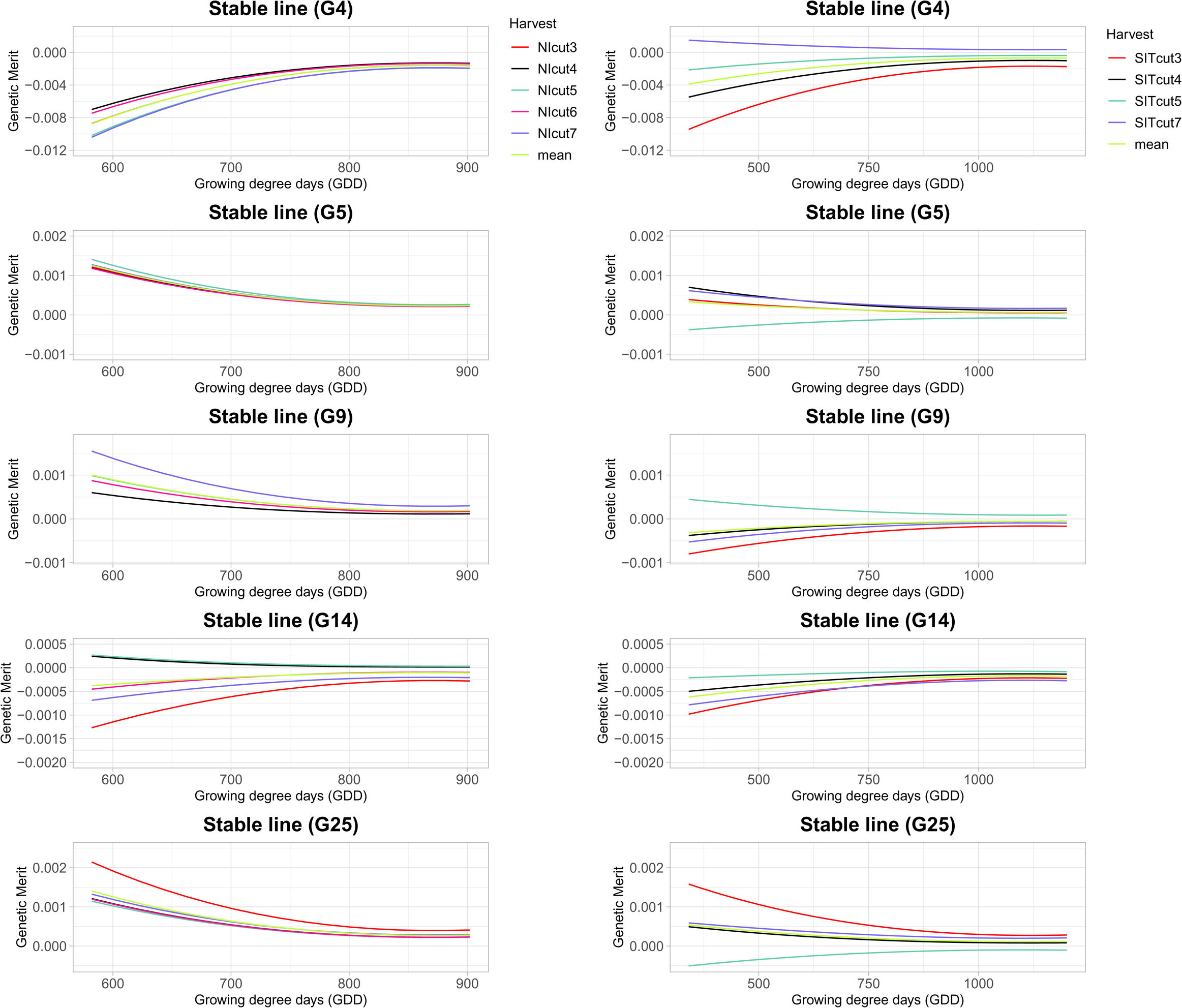
Growth curves derived from GNDVI of five stable alfalfa cultivars across five different harvest seasons of normal irrigation and four different harvest seasons of early termination. The left-hand side figures and right-hand side figures represents growth curves of stable cultivars in normal irrigation condition (NI) and summer irrigation termination condition (SIT) respectively.

**Fig. 19.**
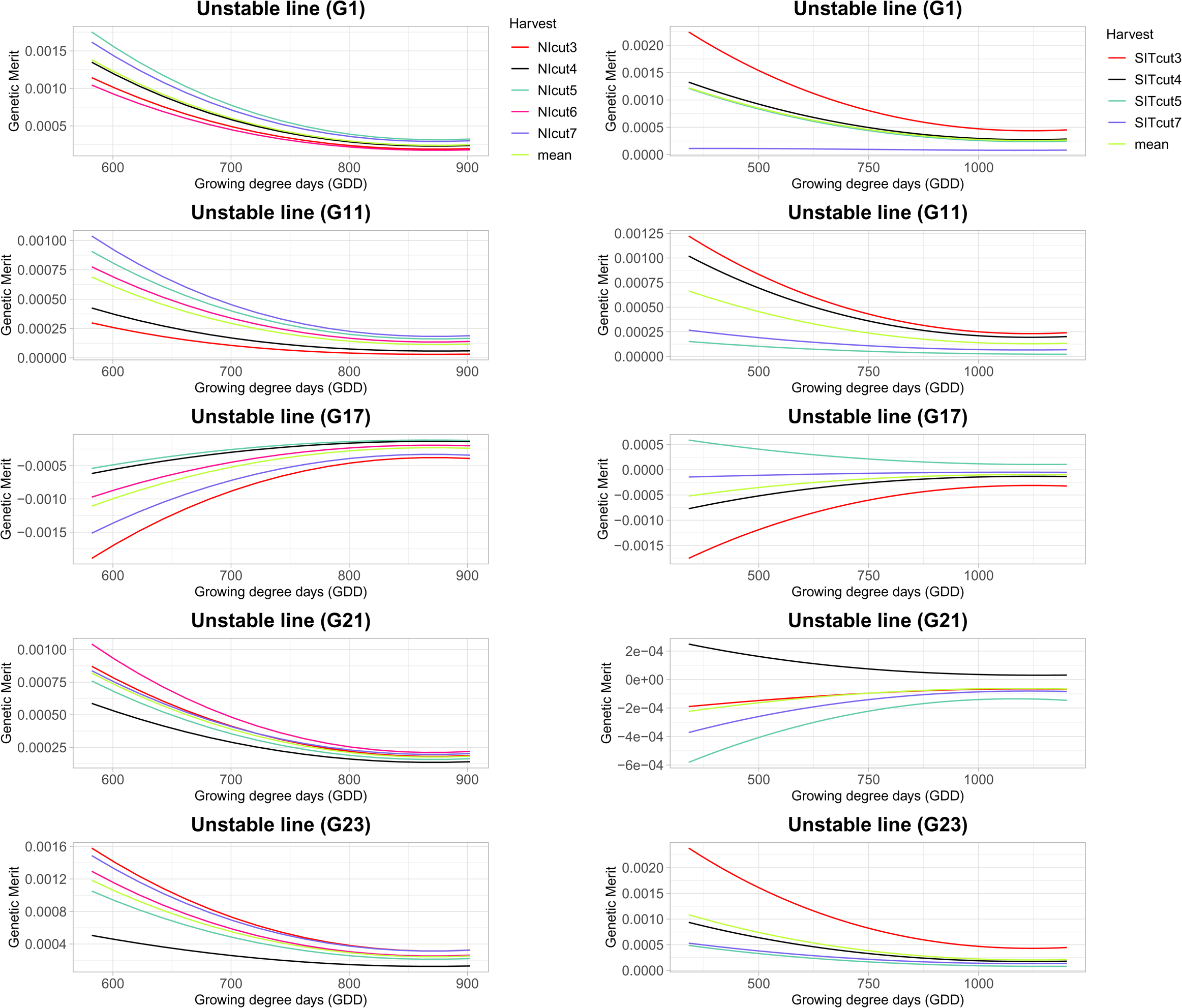
Growth curves derived from GNDVI of five unstable alfalfa cultivars across five different harvest seasons of normal irrigation and four different harvest seasons of early termination. The left-hand side figures and right-hand side figures represents growth curves of stable cultivars in normal irrigation condition (NI) and summer irrigation termination condition (SIT) respectively.

**Fig. 20.**
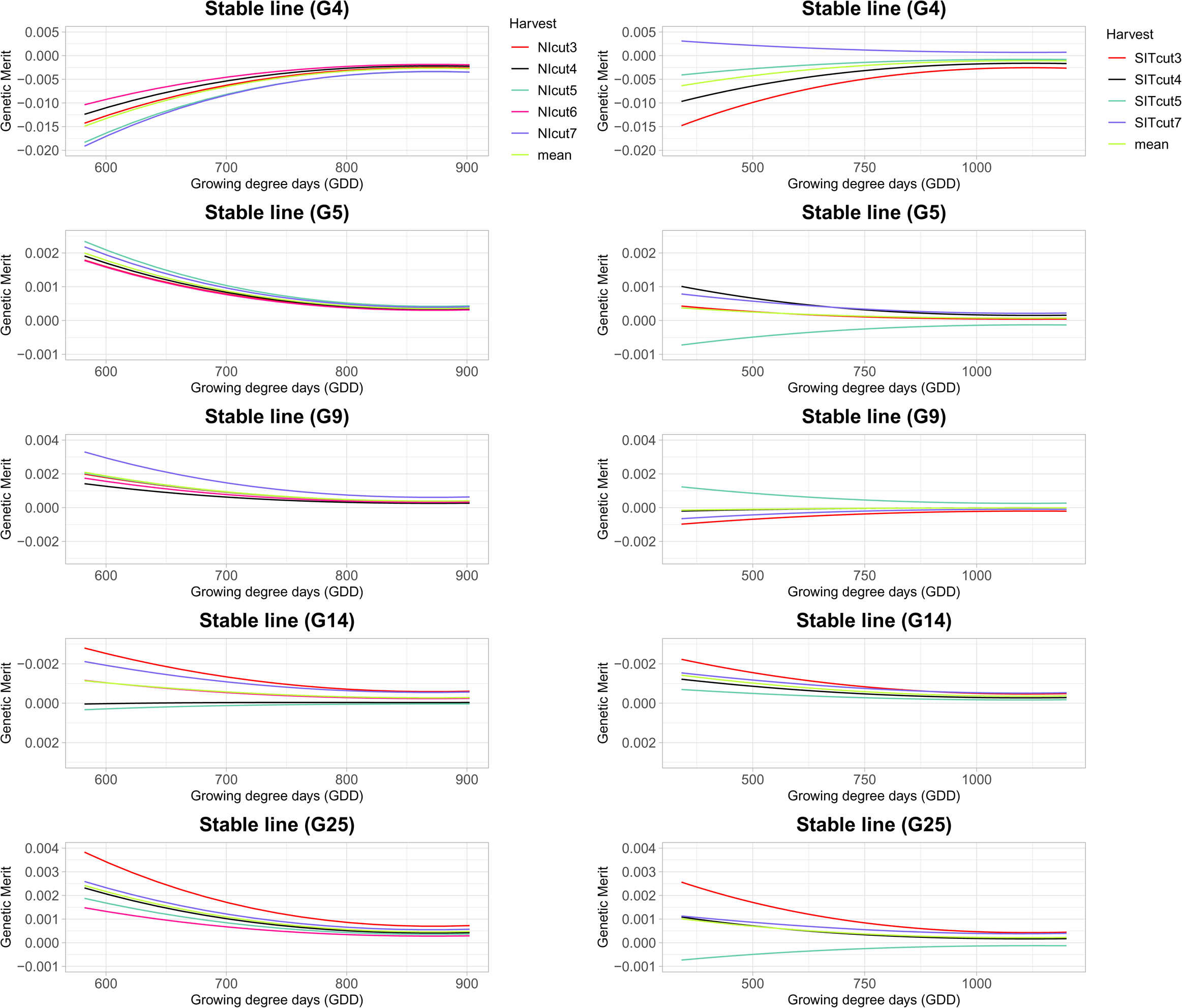
Growth curves derived from NDVI of five stable alfalfa cultivars across five different harvest seasons of normal irrigation and four different harvest seasons of early termination. The left-hand side figures and right-hand side figures represents growth curves of stable cultivars in normal irrigation condition (NI) and summer irrigation termination condition (SIT) respectively.

**Fig. 21.**
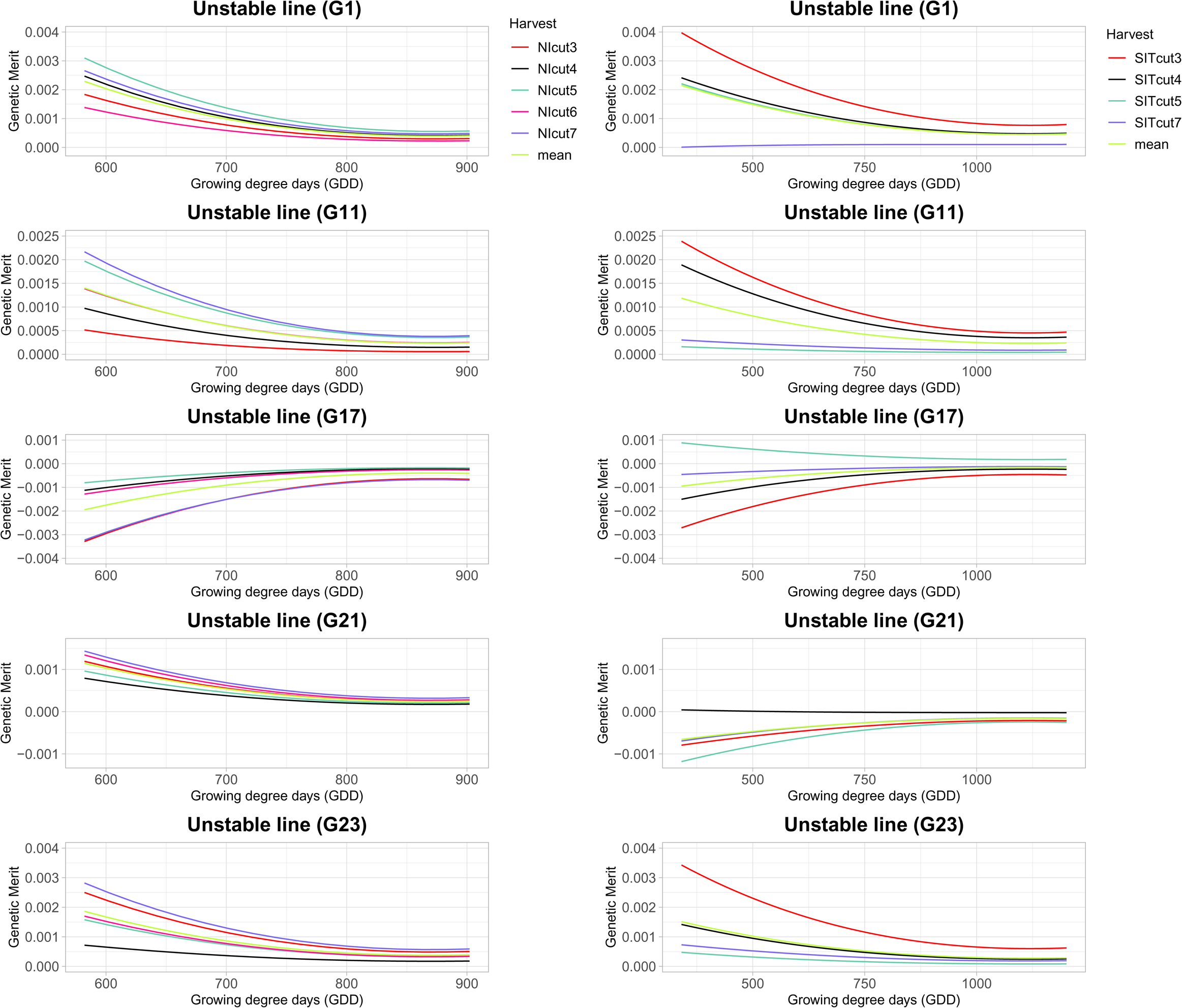
Growth curves derived from NDVI of five unstable alfalfa cultivars across five different harvest seasons of normal irrigation and four different harvest seasons of early termination. The left-hand side figures and right-hand side figures represents growth curves of stable cultivars in normal irrigation condition (NI) and summer irrigation termination condition (SIT) respectively.

**Fig. 22.**
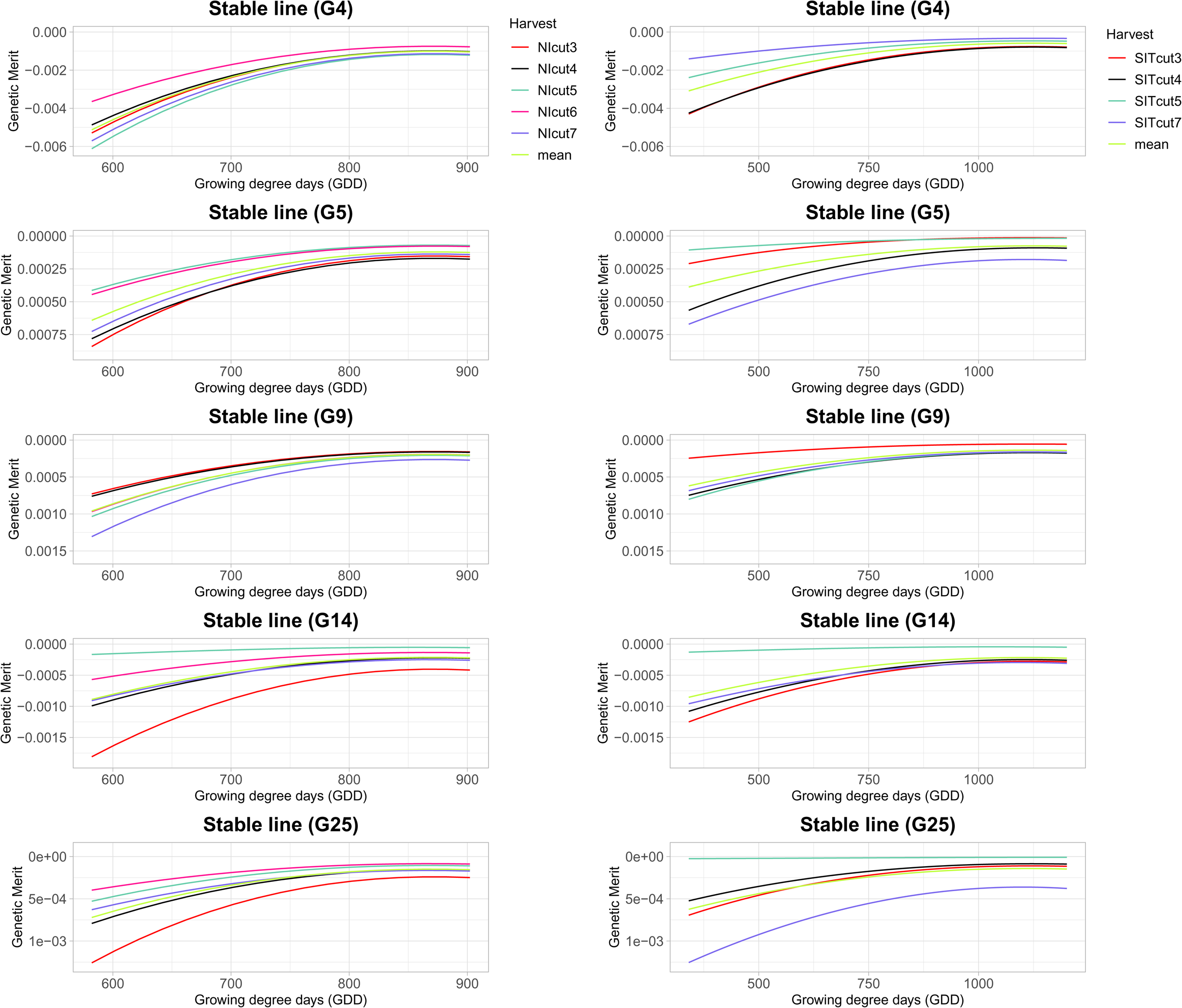
Growth curves derived from NIR of five stable alfalfa cultivars across five different harvest seasons of normal irrigation and four different harvest seasons of early termination. The left-hand side figures and right-hand side figures represents growth curves of stable cultivars in normal irrigation condition (NI) and summer irrigation termination condition (SIT) respectively.

**Fig. 23.**
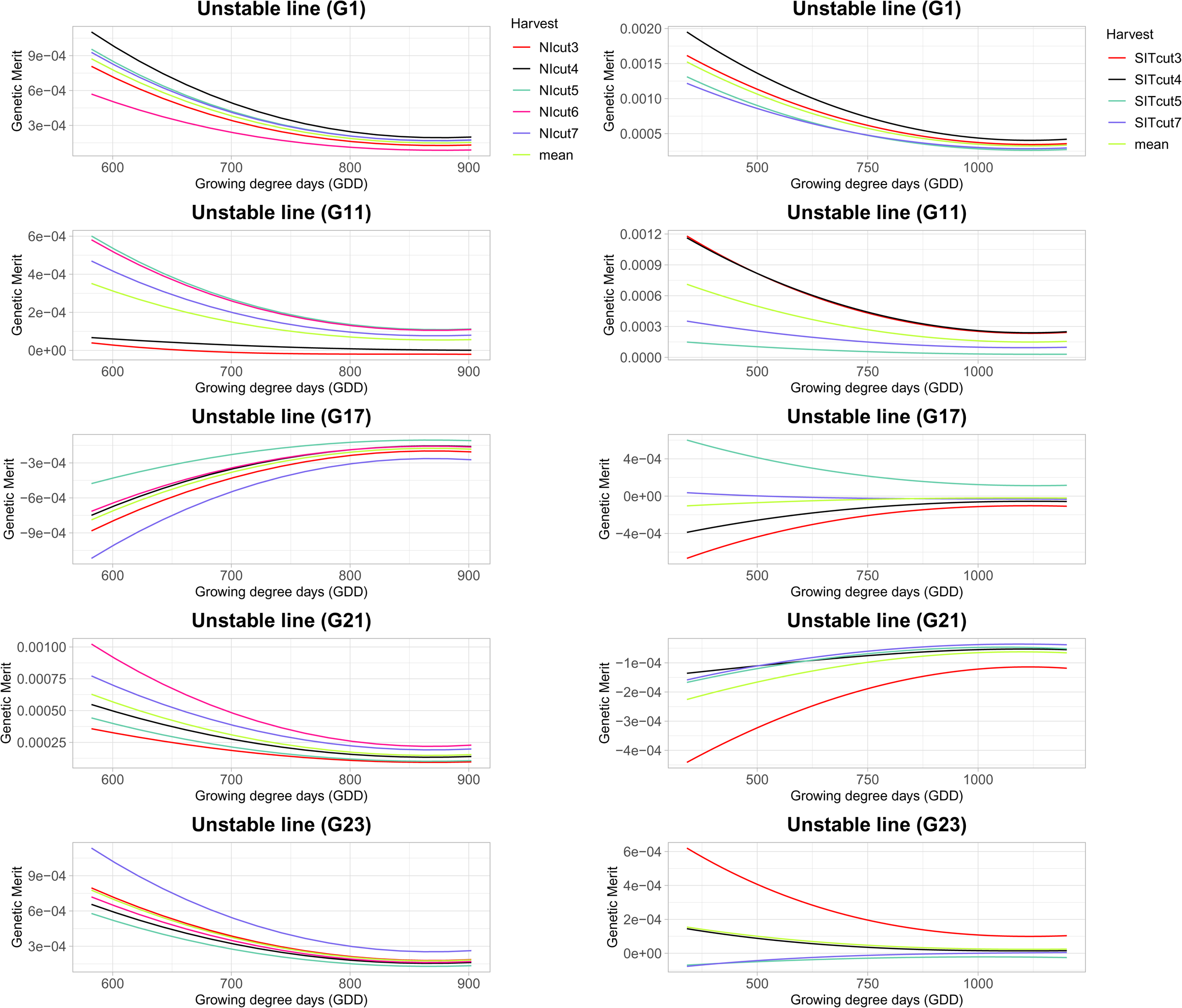
Growth curves derived from NIR of five unstable alfalfa cultivars across five different harvest seasons of normal irrigation and four different harvest seasons of early termination. The left-hand side figures and right-hand side figures represents growth curves of stable cultivars in normal irrigation condition (NI) and summer irrigation termination condition (SIT) respectively.

**Fig. 24.**
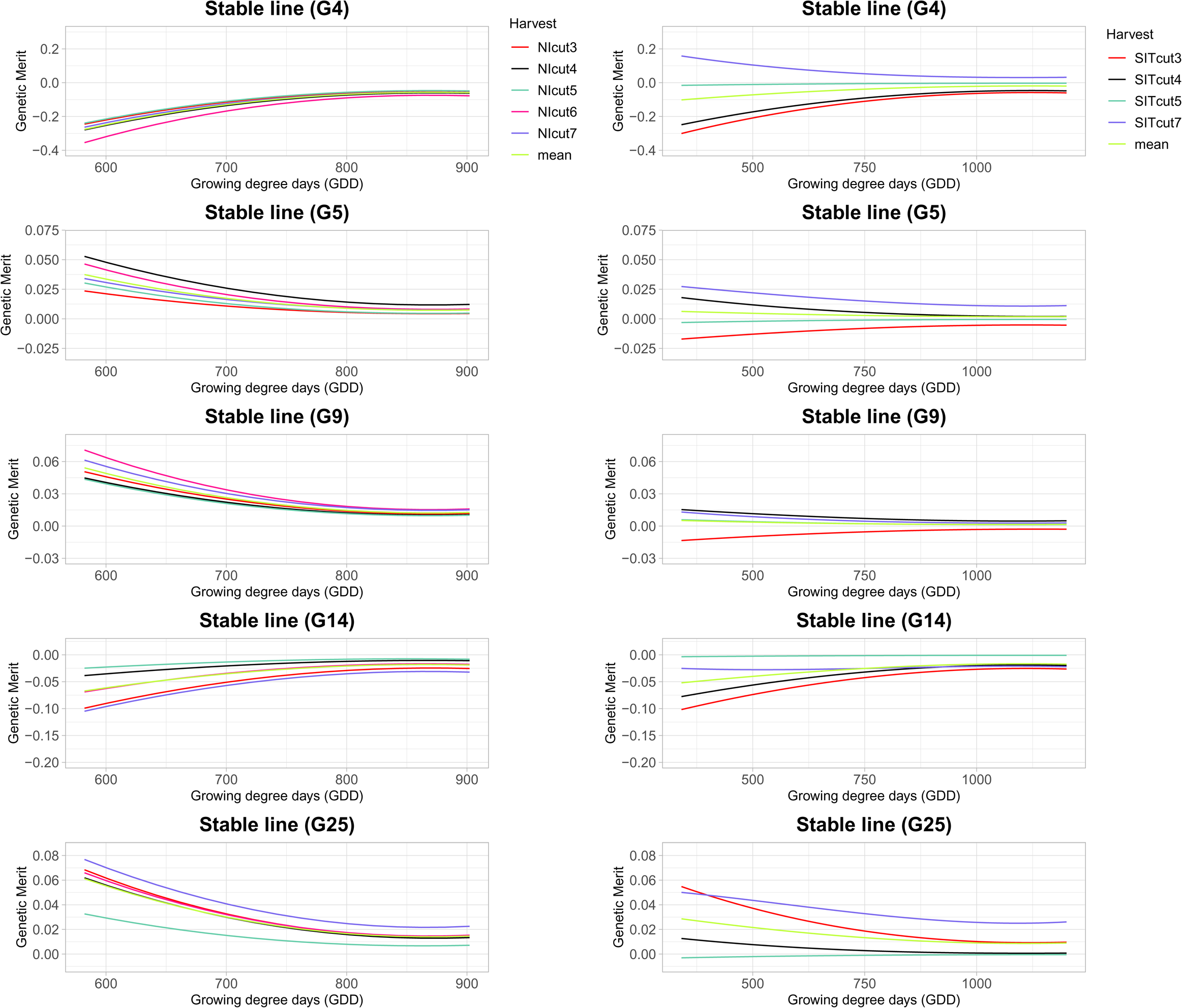
Growth curves derived from Ratio of five stable alfalfa cultivars across five different harvest seasons of normal irrigation and four different harvest seasons of early termination. The left-hand side figures and right-hand side figures represents growth curves of stable cultivars in normal irrigation condition (NI) and summer irrigation termination condition (SIT) respectively.

**Fig. 25.**
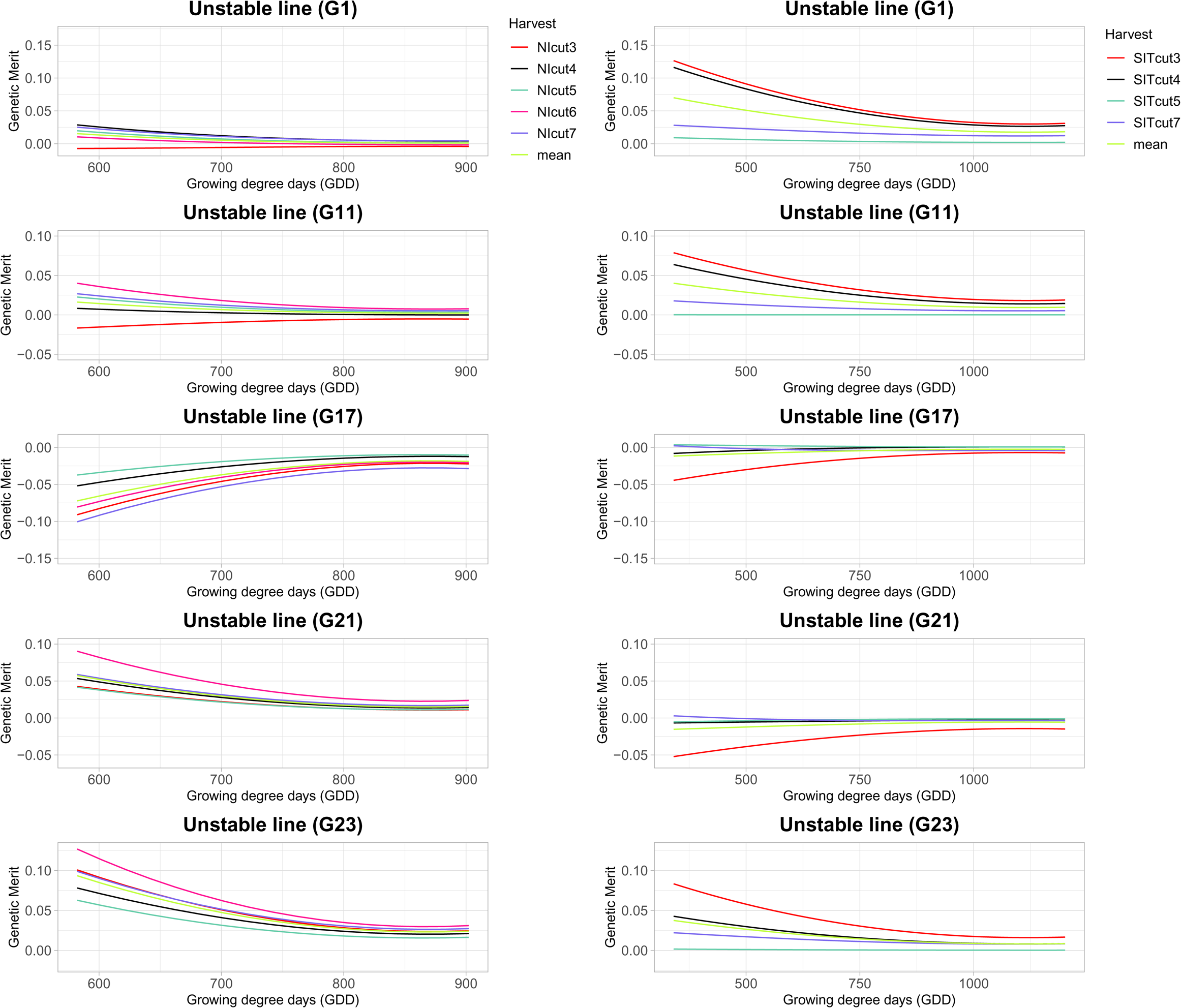
Growth curves derived from Ratio of five unstable alfalfa cultivars across five different harvest seasons of normal irrigation and four different harvest seasons of early termination. The left-hand side figures and right-hand side figures represents growth curves of stable cultivars in normal irrigation condition (NI) and summer irrigation termination condition (SIT) respectively.

### Correlation of variance in yield and variance in genetic merit of phenotypic indices across different environments

The variance in yield of all genotypes across different harvests was calculated for both NY and NMSU trials. Similarly, the variance in genetic merit of all genotypes for VIs at different time points across cuttings was calculated to determine if there was a relationship between variation in growth curves and variation in harvest biomass. The estimated correlation between the variance in yield and variance in genetic merit estimated from VIs at different time points showed a significant correlation in NY, with values ranging from 0.61 to 0.67, 0.63 to 0.66, 0.60 to 0.71, 0.66 to 0.68, and 0.37 to 0.43 for NDVI, GNDVI, NDRE, NIR and Ratio respectively (Table 1). For trials in NM, the correlation between the variance in harvest biomass and the variance in genetic merit estimated from growth curves of all genotypes at different time points across different harvests showed correlations that ranged from 0.19 to 0.35, 0.27 to 0.44, 0.16 to 0.36, 0.68 to 0.79 and 0.91 to 0.93 for GNDVI, NDVI, NDRE, NIR and Ratio, respectively (Table 2).

**Table 1.** Correlation of variance of yield and variance of genetic merit estimated from random regression legendre polynomial (RRLP) model with VIs (GNDVI, NDVI, NDRE, NIR, and Ratio) of all cultivars across different environments of the Ithaca, NY trial at different time points of growing season.

| Growing Degree Days (GDD) | GNDVI | NDVI | NDRE | NIR | Ratio |
| --- | --- | --- | --- | --- | --- |
| 178.55 | 0.65* | 0.67* | 0.6* | 0.68* | 0.43* |
| 218.55 | 0.64* | 0.67* | 0.61* | 0.68* | 0.43* |
| 258.55 | 0.64* | 0.67* | 0.62* | 0.68* | 0.42* |
| 298.55 | 0.63* | 0.67* | 0.64* | 0.67* | 0.42* |
| 338.55 | 0.64* | 0.65* | 0.63* | 0.66* | 0.37 |
| 358.55 | 0.64* | 0.65* | 0.66* | 0.66* | 0.42* |
| 378.55 | 0.65* | 0.65* | 0.67* | 0.66* | 0.42* |
| 418.55 | 0.66* | 0.63* | 0.69* | 0.65* | 0.42* |
| 438.55 | 0.66* | 0.62* | 0.69* | 0.65* | 0.42* |
| 478.55 | 0.65* | 0.61* | 0.71* | 0.66* | 0.42* |

\* indicates P-value < 0.05

**Table 2.** Correlation of variance of yield and variance of genetic merit estimated from random regression legendre polynomial (RRLP) model with VIs (GNDVI, NDVI, NDRE, NIR, and Ratio) of all cultivars across different environments of the NMSU trial at different time points of growing season

| Growing Degree Days (GDD) | GNDVI | NDVI | NDRE | NIR | Ratio |
| --- | --- | --- | --- | --- | --- |
| 342 | 0.19 | 0.27 | 0.16 | 0.68* | 0.91* |
| 402 | 0.2 | 0.28 | 0.18 | 0.69* | 0.91* |
| 462 | 0.22 | 0.3 | 0.2 | 0.7* | 0.91* |
| 522 | 0.24 | 0.33* | 0.22 | 0.72* | 0.92* |
| 582 | 0.26 | 0.35* | 0.25 | 0.73* | 0.92* |
| 642 | 0.29 | 0.38* | 0.28 | 0.75* | 0.92* |
| 702 | 0.31* | 0.4* | 0.31 | 0.76* | 0.92* |
| 762 | 0.33* | 0.42* | 0.34* | 0.77* | 0.93* |
| 822 | 0.35* | 0.42* | 0.35* | 0.78* | 0.93* |
| 902 | 0.35* | 0.44* | 0.36* | 0.79* | 0.93* |

\* indicates P-value < 0.05

## Discussion

One of the objectives of this study was to evaluate the heritability of VIs derived from MSIs and their genetic correlation with the terminal trait biomass yield. Results of this study showed that the VIs have a moderate heritability (Fig 3, Fig 4) comparable to the heritability of harvest biomass. Lower heritability was attributed to poor days of imaging such as the days with cloudy and windy weather. Babar et al. (2007) reported moderate to high heritability of spectral reflectance indices (SRIs) and higher heritability than for grain yield in wheat. Petsoulas et al. (2022) reported moderate to high level of broad sense heritability where the heritability of NDRE ranged from 0.292 to 0.879 and heritability of NDVI ranged from 0.446 to 0.928 in sesame. In the same study, heritability of VIs were reported to be increased with growth stages and started to reduce entering the ripening stage of sesame whereas Anche et al. (2020) reported lower heritability of VIs in early reproductive stage and higher heritability estimates at mid-reproductive stage and late reproductive stage of maize. Another study from (Galán et al. 2020), showed moderate to high heritability estimates (H2 > 0.50) of 23 VIs in winter rye hybrids estimated from hyperspectral reflectance data. Sun et al. (2017) reported that the heritability of NDVI and NDRE ranged from moderate to high across different locations of wheat trial. Sharma et al. (2022) reported consistently higher heritability of VARI and NDVI across growth phases and locations where NDVI and VARI had higher heritability than dry biomass yield. In our study, among all five VIs, GNDVI had highest value of maximum and median heritability. GNDVI measures reflection in near infra-red region and green region of the electromagnetic spectrum (Gitelson et al. 1996). GNDVI provides information about chlorophyll A concentration in plants. The higher heritability of GNDVI might be due to the high biomass of the crop. Sandhu et al. (2021) reported GNDVI as the best predictor of grain protein content of wheat. Previous studies (Hassan et al. 2019; Yang et al. 2020) also reported GNDVI and NDRE as the best predictor of grain yield and nutrient uptake efficiencies across the growth stages.

Multi-trait models were fit to evaluate the correlation between VIs at different time points and harvest biomass yield. The genetic correlations of all five VIs and the biomass yield was found to be strong and statistically significant for all harvests and years. Among five VIs, NIR, NDVI and Ratio had the strongest genetic correlations with biomass. Natarajan et al. (2019) reported a strong correlation between NDVI and sugarcane stalk population and sugarcane yield suggesting that canopy reflectance measurements at an early stage could be used as a screening tool to estimate yield potential. Another study by Prabhakara et al. (2015) used NDVI for prediction of biomass percentage of ground cover in winter forage crops. Other studies have also reported significant association between NDVI and both biomass and GY in irrigated or high-rainfall conditions (Reynolds et al. 1999; Aparicio et al. 2000; Freeman et al. 2003; Gutiérrez-Rodríguez et al. 2004; Babar et al. 2006a; Prasad et al. 2007b; Erdle et al. 2013; Christopher et al. 2014) drought stress (Gutiérrez-Rodríguez et al. 2004; Babar et al. 2006b; Reynolds et al.) and heat stress environments (Reynolds et al. ; Gutierrez et al. 2010; Hazratkulova et al. 2012; Lopes and Reynolds 2012). NDVI was also reported to predict grain yield in soybean (Ma et al. 2001), winter wheat (Raun et al. 2001), and durum wheat (Aparicio et al. 2000). The VIs NDVI, GNDVI, SAVI, G-R were reported to be accurate for estimating biomass at an early stage (Prabhakara et al. 2015) and they were saturated at later stages (Mutanga and Skidmore 2004; Thenkabail et al. 2000). Chen et al. (2009) reported TVI (Triangular Vegetative Index) as useful index for predicting canopy biomass at later stage. NDVI and SR are based on the red (visible) and NIR wavelengths and give higher values at early growth stages, but their values decrease with the advancement in growth cycle because plants are losing photosynthetically active plant parts. Serrano et al. (2000) reported that simple ratio (SR) can reliably predict winter wheat grain yield under nitrogen stresses. Among the three spectral indices, simple ratio (SR), normalized difference vegetation index (NDVI), and photochemical reflectance index (PRI), SR was identified as the best index for assessment of crop growth and yield in durum wheat (Aparicio et al., 2000). Another study by Gutierrez et al. (2004) found the strongest correlation of SR and NIR with cotton lint yield showing 60% and 58% of variations in cotton lint yield respectively. In the same study, SR and NIR had higher coefficients of determination in cotton biomass and leaf area index (LAI) compared to NDVI as these indices were not saturated at late growth stage whereas (Aparicio et al. 2000; Aparicio et al. 2002) reported that NDVI and SR were not able to predict variations in biomass successfully when estimated at later growth stages of durum wheat. Hence, the use of multiple indices is recommended to get better predictions of biomass yield as different types of VIs are sensitive to different stages of crop growth and amount of biomass. The high heritability and strong genetic correlation between VIs and biomass yield of alfalfa in our study suggest that VIs can be used as a selection tool and help plant breeders to reliably evaluate cultivars in a fast and nondestructive (Lobos et al. 2019; El-Hendawy et al. 2019; Prasad et al. 2007a; Babar et al. 2007; Gutierrez et al. 2010).

RR models with third order Legendre polynomials provided the best fit and were used to model the growth curve trajectories using VIs as phenotypes. Estimate RR coefficients were used to obtain breeding values (BVs) for all time points between the first day of imaging and harvest. Sun et al (2017) used RR model with cubic splines in wheat (*Triticum aestivum*) to obtain best linear unbiased predictions of secondary traits derived from high-throughput hyperspectral and thermal imaging. RR model with a linear spline was also reported as a potential alternative approach to mixed model to fit the VIs from multiple time points (Anche et al. 2020), but Legendre polynomials were found to provide a better fit to maize data in subsequent analyses (Anche et al. 2023). When cumulative indices were used as phenotypes, the correlation was found to increase through time (Fig 3-6). This could be because cumulative indices accounts for earlier season VI data, and therefore becomes more informative than raw data on predicting biomass yield of the growing season. Similar results were reported in maize (Anche et al. 2023), concluding that cumulative VIs were better phenotype to model the covariance structures as they provided more stable and consistent results compared to using raw VIs as a phenotype.

In our study, we observed a decreasing trend in the variance components over time for each harvest. Higher genetic variation was observed in the breeding values of VIs in early growth stages compared to later stages as cultivars reached full canopy cover. In alfalfa stands, allowing the crop to reach maximum vegetation saturation before flowering is the ideal balance to develop maximum biomass while also maintaining nutritional quality. A declining ability of spectral indices to discriminate different genotypes was reported in other crops as the canopy closes and its spectral reflectance saturated (Marti et al. 2007). In this study, all VIs showed strong correlations with biomass yield across all time points, and the growth trajectories could separate high yielding and low yielding genotypes rapidly and efficiently starting in the early stage of growth season.

The moderate heritability and moderate to strong genetic correlations with harvest biomass observed in NY and NM trials, indicate that VIs collected via UAV can be used to model temporal genetic variation associated with harvest biomass yield. RR models provided a parsimonious approach to estimate temporal covariance functions and assess cultivar persistence and stability, which can be affected by biotic and abiotic effects of the environment. The RR model depicted dynamic aspects of phenotypes, which can enable better analysis cultivar plasticity, adaptability, stability and yield performance (Alves et al. 2020) across a range of dynamic environmental conditions through growth periods. As such, information on growth curves can provide additional information for selecting lines that are best adapted to the target environments.

The growth trajectories of stable genotypes and unstable genotypes in NI and SIT termination of NMSU trial showed more instability in growth curves in SIT than NI (Figs 18 to 25). This is likely due to higher genetic variance among different cultivars in stressed environment compared to normal condition and indicates that growth parameters may provide additional information on stress tolerance. The observed correspondence in plasticity of growth curves and stability in biomass harvest demonstrate the potential to model GxE temporally throughout the growth period as a function of dynamic environmental variables. Esten et al. (2018) reported stronger correlation of NDVI and GY (r = 0.25 – 0.54) and NDVI and biomass (r = 0.17 – 0.46) in lowest yielding sites-years. In the same study, NDVI was reported to have greater ability to detect biomass differences between lines in low-yielding environments, where canopy closure was not present. Similar results were previously presented where stronger correlations of NDVI and grain yield was observed under abiotic stress compared with high-yielding environments (Gutiérrez-Rodríguez et al. 2004; Gutierrez et al. 2010; Lopes and Reynolds 2012).

## Conclusion

The use of multi-spectral imaging for alfalfa over the growing seasons in NY and NMSU demonstrated that VIs are heritable and that genetic correlations were significant for most time points and years. The measurement of cumulative NDVI showed that correlations of NDVI to biomass increased over time closer to harvest/cutting date. Strong correlations of NDVI to biomass harvest increase the possibility of using MSI to reduce the amount of biomass harvest phenotyping needed, potentially reducing phenotyping costs. The use of random regressions and Legendre polynomials demonstrated that longitudinal modeling of VIs can capture genetic variation, and stability in growth curves across cuttings was associated with stability in harvest biomass over harvests, years, locations and irrigation treatments. These results indicate that random regressions of VIs captures throughout a growth period can provide a greater dynamic understanding of aspects of phenotypic plasticity, stability and yield performance for crop improvement.

## Data availability

R Code and data are available in the github: https://github.com/rthapa1/FFAR_RandomRegressionModel_growthcurve_modelling_stabilitya nalysis_alfalfa

## Authors contributions

Ranjita Thapa: Processed MSI data; Developed statistical model; Conducted statistical analysis; Investigation; Methodology; Software; Validation; Visualization; Writing – original draft, review & editing. Karl H. Kunze: Review & editing. Julie Hansen: Review & editing. Christopher Pierce: Data collection; Review & editing. Virginia Moore: Review & editing. Ian Ray: Funding acquisition; Validation; Visualization; review & editing. Liam Wickes-Do: Data collection. Nicolas Morales: Data collection. Felipe Sabadin: Review & editing. Nicholas Santantonio: Conceptualization; Funding acquisition; Writing – review & editing. Michael A Gore: Funding acquisition; Writing – review & editing. Kelly Robbins: Conceptualization; Funding acquisition; Methodology; Project administration; Resources; Supervision; Validation; Visualization; review & editing

## Acknowledgments

We would like to thank Jesse Chavez, Ryan Crawford and Jamie Crawford for their assistance with harvesting and data collection of the NY alfalfa trials.

## Abbreviations

ATC: average tester co-ordination
RCBD: randomized complete block design
BLUE: best linear unbiased estimators
FY: Forage yield
GDD: growing degree days
GGE: genotype main effects and genotype by environment
GNDVI: green normalized difference vegetation index
G×E: genotype by environment
HTP: high-throughput phenotyping
MSI: multispectral imaging
NDRE: normalized difference red edge index
NDVI: normalized difference vegetation index
OSAVI: optimized soil adjusted vegetation index
NIR: near infrared
NMSU: New Mexico State University
RR: random regression
RRLP: random regression Legendre polynomial
SCCCI: simplified canopy chlorophyll content index
ST-GBLUP: single trait genomic linear unbiased prediction
VARI: visible atmospherically resistant index
VI: vegetative index
cVI: cumulative vegetative index
UAV: unmanned aerial vehicle

